# Human APOBEC3B promotes tumor heterogeneity *in vivo* including signature mutations and metastases

**DOI:** 10.1101/2023.02.24.529970

**Authors:** Cameron Durfee, Nuri Alpay Temiz, Rena Levin-Klein, Prokopios P. Argyris, Lene Alsøe, Sergio Carracedo, Alicia Alonso de la Vega, Joshua Proehl, Anna M. Holzhauer, Zachary J. Seeman, Yu-Hsiu T. Lin, Rachel I. Vogel, Rocio Sotillo, Hilde Nilsen, Reuben S. Harris

## Abstract

The antiviral DNA cytosine deaminase APOBEC3B has been implicated as a source of mutation in many different cancers. Despite over 10 years of work, a causal relationship has yet to be established between APOBEC3B and any stage of carcinogenesis. Here we report a murine model that expresses tumor-like levels of human APOBEC3B after Cre-mediated recombination. Animals appear to develop normally with full-body expression of APOBEC3B. However, adult males manifest infertility and older animals of both sexes show accelerated rates of tumorigenesis (mostly lymphomas or hepatocellular carcinomas). Interestingly, primary tumors also show overt heterogeneity, and a subset spreads to secondary sites. Both primary and metastatic tumors exhibit increased frequencies of C-to-T mutations in TC dinucleotide motifs consistent with the established biochemical activity of APOBEC3B. Elevated levels of structural variation and insertion-deletion mutations also accumulate in these tumors. Together, these studies provide the first cause-and-effect demonstration that human APOBEC3B is an oncoprotein capable of causing a wide range of genetic changes and driving tumor formation *in vivo*.

## INTRODUCTION

Cancer development and progression are evolutionary processes driven by mutations and further fueled by epigenetic alterations and environmental factors (reviewed by refs.^1–3^). Major advances over the past decade in genome sequencing and computational technologies have provided an unprecedented view of the entire landscape of genomic alterations that occurs in cancer. This has enabled precise documentation of the many oncoproteins and tumor suppressors that contribute to over 50 different human cancer types. Another profound advance enabled by these technologies is a capacity to extract distinct mutation signatures from otherwise extremely complex montages of mutational events in single tumors (reviewed by refs.^4–6^). Upon extension to large numbers of tumors, the abundance of each distinct signature becomes starkly apparent and, taken together with chemical, biological, and genetic information, informs guesses as to the most likely etiologic source (endogenous or exogenous) of the DNA damage that led to the observed signature. A few of many robust examples to-date include spontaneous, water-mediated deamination of methyl-C to T in CG motifs (COSMIC single base substitution signature 1, SBS1), C-to-T mutations in di-pyrimidine motifs caused by A-insertion opposite UV light-catalyzed pyrimidine dimers (SBS7), and APOBEC-catalyzed C-to-U deamination events in TC motifs leading to C-to-T and C-to-G mutations (SBS2 and SBS13, respectively).^5^

The human APOBEC family of polynucleotide C-to-U deaminase enzymes is comprised of apolipoprotein B mRNA editing catalytic subunit 1 (the family namesake APOBEC1), activation-induced cytidine deaminase (AICDA but popularly called AID), and seven distinct APOBEC3 enzymes (A3A, B, C, D, F, G, and H; reviewed by refs.^7–9^). APOBEC1 functions in mRNA editing, AID in antibody gene diversification, and A3A-H in virus restriction. Although most of these enzymes preferentially deaminate TC motifs in single-stranded (ss)DNA, a number of studies have converged on A3A and A3B as the major sources of APOBEC signature mutations in cancer (refs.^10, 11^ and references therein). Specifically, expression of A3A or A3B triggers an abundance of APOBEC signature mutations in human cells, and CRISPR-mediated gene knockouts lower the capacity of cancer cell lines to accumulate both SBS2 and SBS13 mutation signatures. Summaries of relevant literature including clinical correlations have been published recently (reviewed by refs.^12, 13^).

A major obstacle in assessing the overall impact of A3A and A3B in cancer is a lack of appropriate murine models. Mice encode homologs of human APOBEC1 and AID but lack direct equivalents of human A3A and A3B (*i.e*., mice encode only a single Apobec3 protein with a domain organization not found in humans). Moreover, murine Apobec3 is cytoplasmic, and APOBEC signature mutations as defined above do not occur naturally in mice (reviewed by refs.^14^). However, recent studies have begun to overcome this obstacle by developing murine model systems to study mutagenesis by human A3A and A3B. First, a transgenic line that expresses low levels of human A3A has no cancer phenotypes alone but is capable of enhancing the penetrance of Apc^Min^-driven colorectal tumors and causing an accumulation of SBS2 (but not SBS13) signature mutations.^15^ Second, hydrodynamic delivery of human A3A into murine hepatocytes, coupled to liver regeneration by selecting for Fah function, results in hepatocellular carcinoma development within 6 months.^15^ Both SBS2 and SBS13 mutation signatures are evident in these liver tumors but expression of human A3A is selected against and lost, which limits the potential for longer-term studies on tumor evolution. Moreover, A3B expression is aphenotypic over the same duration in this system. Last, low levels of human A3B expressed constitutively from the endogenous *Rosa26* promotor cause no overt tumor phenotypes and no detectable APOBEC signature mutations.^16^ The latter two studies call-to-question the role of A3B in human tumor pathology.

Thus, it is not yet known if human A3B is capable of driving oncogenesis *in vivo* or if the mutations it is implicated in causing are simply passenger events in cancer or maybe even caused by another A3 family member such as A3A. To address these and other questions we have created a new murine model for inducible expression of human A3B. In these animals, a human *A3B* minigene is integrated into the *Rosa26* locus downstream of the *Rosa26* promoter, a stronger heterologous CAG promoter, and a strong transcription stop cassette flanked by *loxP* sites. Therefore, Cre-mediated removal of the transcription stop cassette results in strong A3B expression. This model enabled us to show that full-body expression of CAG-driven levels of human A3B recapitulates protein amounts reported in many human cancers. Young animals show no overt phenotypes except that males never become fertile. In comparison to naturally aged wildtype animals, CAG-A3B mice of both sexes develop tumors, predominantly blood and liver cancers, an average of 5.2 months earlier. A subset of animals also show clear evidence for metastasis. Both primary and metastatic tumors manifest a clear APOBEC mutation signature (SBS2), which interestingly also associates with an elevated occurrence of structural variations, including small insertion and deletion mutations (indels) and larger-scale events. Overall, these studies demonstrate that A3B is capable of driving tumor formation and thus provide a new system for studying tumor evolution and undertaking preclinical studies.

## RESULTS

### Construction of a murine model for inducible expression of human A3B

To test the idea that the lack of tumor phenotypes in our original *R26-A3B* model^16^ may be due to low expression levels, we established a new C57BL/6 mouse model for inducible expression of high levels of human A3B by inserting a strong CAG promoter upstream of the transcription stop cassette (*CAG-A3B* versus *R26-A3B* schematics in **Figure 1A**; additional details in **Figure S1**). As anticipated, crossing these mice with CMV-Cre animals to remove the stop cassette and generate progeny with full-body A3B expression (hereafter referred to as *CAG-A3B* mice) results in at least 5-fold higher human A3B protein levels in all tissues examined including liver, pancreas, and spleen (**Figure 1B**). Higher A3B expression also results in elevated ssDNA deaminase activity in the same tissues, with a caveat that activity in splenic extracts is challenging to quantify due to non-specific substrate cleavage by an endogenous nuclease (**Figure 1C**). A3B protein expression is further demonstrated by immunohistochemistry (IHC) with the rabbit monoclonal antibody 5210-87-13 (**Figure 1D**). IHC clearly shows a strong accumulation of human A3B in the nuclear compartment of cells in multiple murine tissues, consistent with prior reports for human A3B subcellular localization in human cell lines and tissues.^17–20^ These observations demonstrate that higher levels of human A3B are tolerated in primary mouse tissues and, further, that its nuclear import mechanism is conserved, despite the fact that only distantly related polynucleotide deaminase family members are expressed in mice (*i.e*., Apobec3, Apobec1, and Aicda).

**Figure 1.**
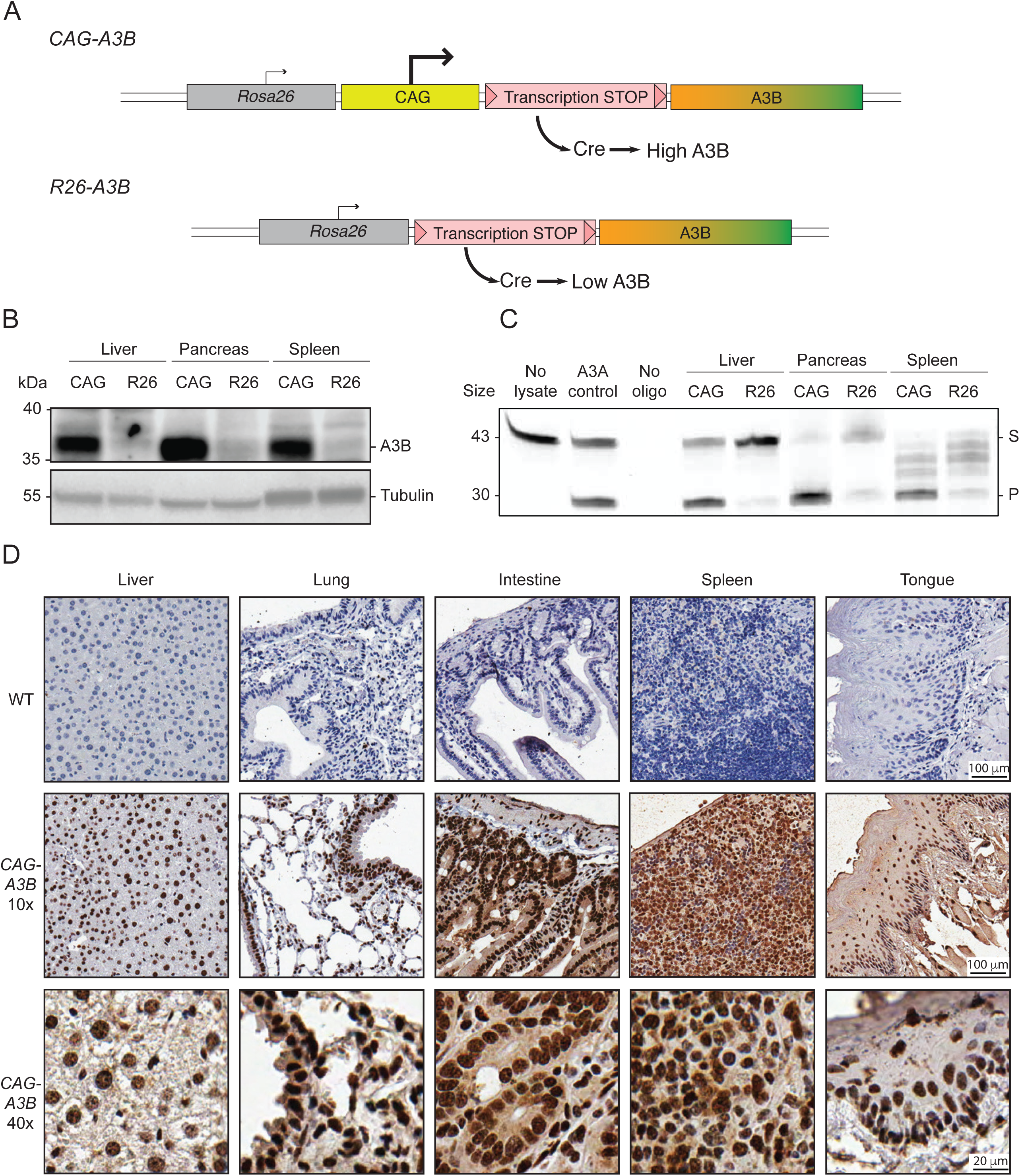
Murine models for inducible expression of low or high levels of human A3B. (**A**) Schematics of *CAG-A3B* and *R26-A3B* knock-in alleles. Human *A3B* expression at high and low levels, respectively, only occurs after Cre-mediated excision of the STOP cassette. (**B-C**) Immunoblot and ssDNA deaminase activity of human A3B protein expressed in the indicated tissues from *CAG-A3B* and *R26-A3B* animals. Tubulin provides a loading control, and recombinant A3A is a positive control for activity, respectively (S, substrate; P, product). (**D**) Anti-A3B IHC staining of representative tissues from WT and *CAG-A3B* mice (40x magnifications are enlargements of regions of the corresponding 10x images). See also **Figure S1**.

### High A3B levels cause male-specific infertility

A striking and unexpected phenotype of *CAG-A3B* animals is male infertility. This is illustrated by no progeny from *CAG-A3B* male x WT female crosses, in comparison to standard Mendelian ratios from *CAG-A3B* female x WT male crosses (**Figure 2A**). In contrast, *R26-A3B* crosses yielded near-Mendelian rations regardless of the sex of the parental animal. Moreover, *R26-A3B* x *R26-A3B* crosses also yield expected numbers of all progeny combinations indicating that 2-fold more *R26-A3B* levels are insufficient to account for the infertility phenotype observed with *CAG-A3B* males.

**Figure 2.**
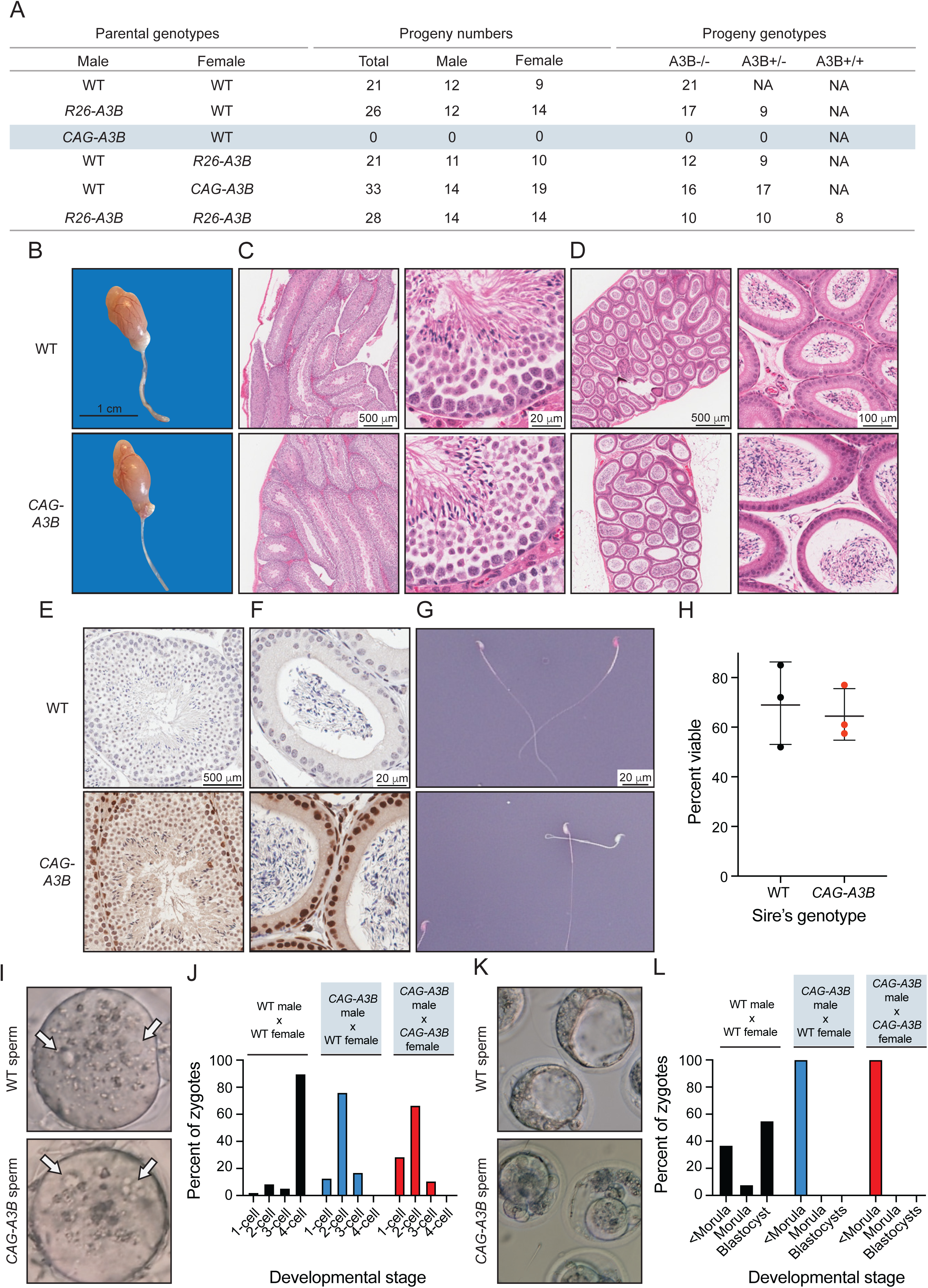
High A3B levels cause male-specific infertility. **(A)** Progeny numbers and genotypes for the indicated crosses (n=3 litters per cross). **(B)** Images of a representative testicle and epididymis from WT and *CAG-A3B* animals. (**C-D**) H&E-stained sections of WT (top) and *CAG-A3B* (bottom) testicle and epididymis, respectively. (**E-F**) Anti-A3B IHC staining of the seminiferous tubule and epididymal lumen from WT and *CAG-A3B* males, respectively. (**G-H**) Representative images and quantification of spermatozoa from WT and *CAG-A3B* males, stained with eosin and nigrosin to distinguish live (white) and dead (pink) cells, respectively (mean +/-SD of n=200 sperm from 3 independent males). (**I**) Images of zygotes 7 hours post-fertilization of a WT ovum with spermatozoa from the indicated male genotypes. Arrows point to pronuclei, indicating fertilization. (**J**) Proportion of embryos at the indicated developmental stage 24 hours post-fertilization *in vitro* (n>50 zygotes analyzed per condition). (**K**) Images of developing embryos 96 hours post-fertilization *in vitro*. (**L**) Proportion of embryos at the indicated developmental stage 96 hours post-fertilization *in vitro* (n>50 zygotes analyzed per condition; continuation of experiment reported in panel J).

Testes from *CAG-A3B* males are morphologically normal at macroscopic and microscopic levels by hematoxylin & eosin (H&E) staining (**Figure 2B**). Seminiferous tubules and epididymal lumen are also normal by H&E staining (**Figure 2C-D**). Moreover, high magnification images of seminiferous tubules and epididymal lumen stain positive for human A3B and show no obvious morphological differences (**Figure 2E-F**). Notably, A3B is localized to the nuclear compartment of germ stem cells and early-stage sperm cells but seems undetectable in the late stages of sperm development including in spermatozoa (**Figure 2E**). Cells within the epithelium of the epididymal lumen also express nuclear A3B, but the adjacent mature spermatozoa appear negative (**Figure 2F**). Moreover, mature sperm from *CAG-A3B* males appear morphologically normal with characteristic hook-shaped heads and functional tails of normal length (**Figure 2G** and **Video S1**). Additionally, eosin & nigrosin staining indicates no significant difference in the number of live sperm from WT versus *CAG-A3B* males (**Figures 2G-H**).

Next, WT female eggs were fertilized *in vitro* with sperm from *CAG-A3B* males and from WT males as controls. In all instances, sperm cells are able to fertilize eggs as evidenced by the appearance of two pronuclei per ovum (**Figure 2I**). However, overt defects become apparent within 24 hours, with all *CAG-A3B* embryos arresting before the 4-cell stage (**Figure 2J**). Moreover, at 96 hours post-fertilization, differences are even more stark with all CAG-A3B embryos visibly terminated (<morula stage development in **Figures 2K-L**). In contrast, WT embryos show normal developmental trajectories (**Figures 2I-L**). These observations combine to suggest that the genetic integrity of *CAG-A3B* male sperm may be compromised.

### *CAG-A3B* mice exhibit accelerated rates of tumor progression and elevated tumor numbers

Our recent studies found no difference in longevity or rates of tumor formation between WT and *R26-A3B* animals, which express *low Rosa26* levels of A3B in most tissues.^16^ This analysis has been expanded here at two different animal facilities (Minneapolis and Oslo) and, again, no significant difference in mouse development or overall rates of tumor formation are observed (**Figures 3A** **and S2A**). However, a slight increase in lymphoma frequency may be apparent in the Oslo facility, where animals house in a minimal disease unit (**Figure S2B**), but this modest phenotype is not accompanied by elevated mutation loads or an obvious APOBEC mutation signature (**Figures S2C-E**). An independent model, in which human *A3B* (as a *turbo-GFP* fusion) is integrated at the *ColA1* locus and expressed inducibly using a *R26*-integrated tetracycline transactivator,^21^ also yields modest A3B expression levels and no significant tumor phenotypes (**Figure S3**).

In contrast to these models that directly or indirectly express low levels of human A3B, our new *CAG-A3B* model with full-body A3B expression shows accelerated rates of tumor formation (**Figure 3A**). By 600 days, over 50% of *CAG-A3B* animals have developed tumors, whereas less than 20% of WT mice are penetrant at this early timepoint (**Figure 3A**). *CAG-A3B* animals also have significantly higher tumor burdens as compared to WT mice consistent with accelerated levels of mutagenesis (**Figure 3B**). Most of the tumors in *CAG-A3B* animals are lymphomas or hepatocellular carcinomas (HCCs) (**Figure 3C**). WT animals also show a similar spectrum of tumors (albeit with longer latencies) suggesting that A3B may accelerate the penetrance of pre-existing cancer predispositions (**Figure 3C**). In support of this possibility, MMTV-Cre is known to have leaky expression in hematopoietic cells^22–24^ and, accordingly, our attempts to induce CAG-A3B specifically in mammary epithelial cells also trigger the formation of lymphomas (**Figure S4**). Importantly, the levels of human A3B expressed in these murine tumors approximate the upper level of those reported in human cancers of multiple different tissue types (**Figure 3D**).

**Figure 3.**
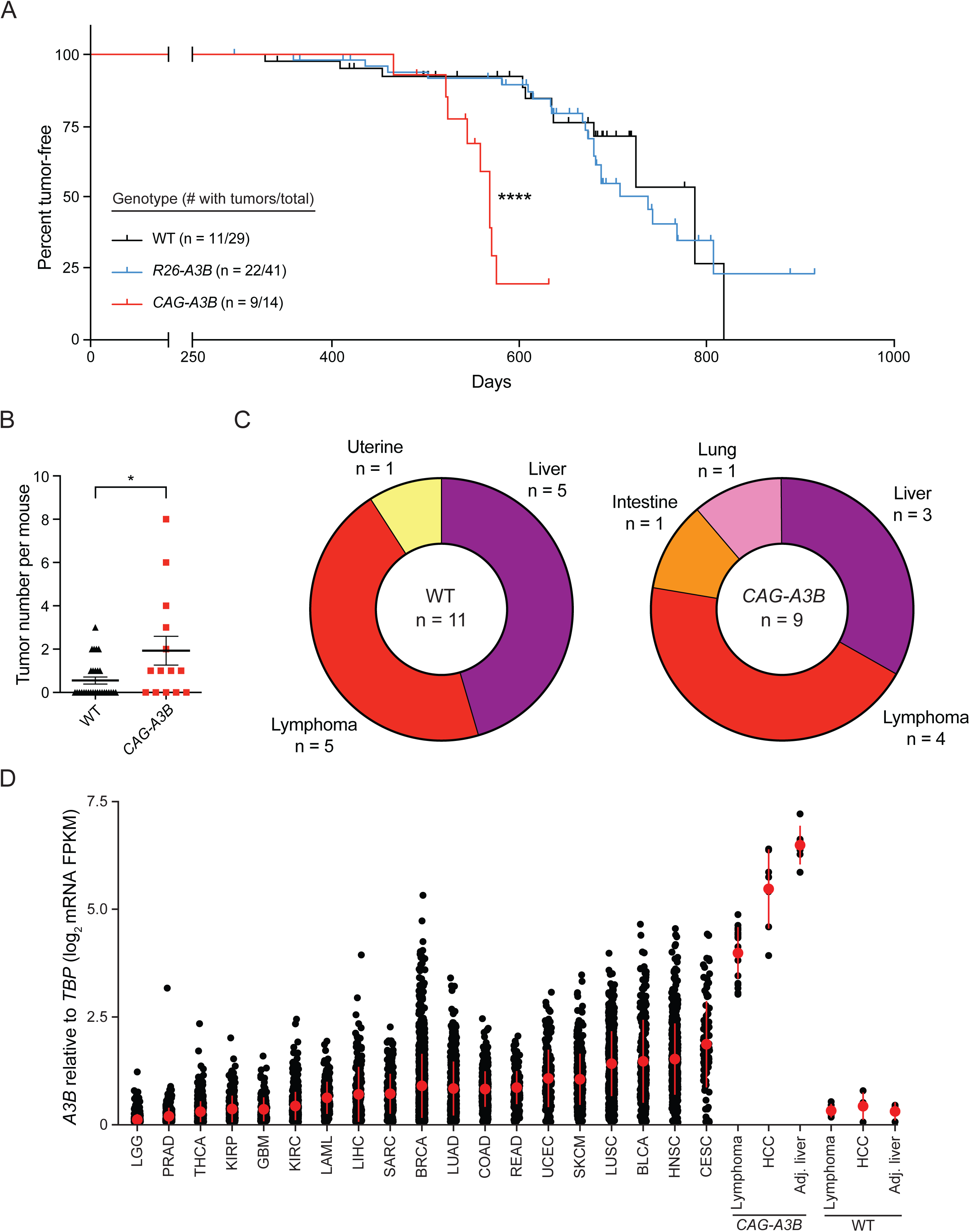
*CAG-A3B* mice exhibit accelerated rates of tumor progression and elevated tumor numbers. **(A)** Kaplan-Meier curves comparing tumor-free survival of WT (n=29), *R26-A3B* (n=41), and *CAG-A3B* (n=14) mice (****, p<0.0001 by log-rank Mantel-Cox test). The number of animals with tumors is shown over the total number of animals in each group. **(B)** Dot plot of the number of tumors per mouse in each respective genotype (mean +/-SEM; *, p=0.0387 by Mann-Whitney U test). **(C)** Pie chart summarizing primary tumor locations in WT and *CAG-A3B* mice. **(D)** *A3B* mRNA expression levels relative to those of the housekeeping gene *TBP* for the indicated specimens (single data points show individual values and the median +/-SD is indicated in red for each group). Human tumor RNA-seq data sets are from TCGA, and WT and *CAG-A3B* data sets are from the indicated tumor and matched normal liver tissues described here (WT lymphoma n=4, HCC n=4, and normal liver n=4; *CAG-A3B* lymphoma n=18, HCC n=7, and normal liver n=6). See also **Figures S2**, **S3**, **S4**, and Table S1.

### Heterogeneity and evidence for metastasis in tumors from *CAG-A3B* animals

Tumors that develop in *CAG-A3B* animals are visibly heterogeneous, which is a hallmark of human tumor pathology that has been difficult to recapitulate in mice (**Figure 4A-D**). For instance, in comparison to normal intestine-associated lymphoid follicles in **Figure 4A** and a normal liver in **Figure 4B**, both lymphomas and HCCs show significant visible heterogeneity (pictures of representative tumors in **Figure 4C-D**; summary of all *CAG-A3B* tumor information in Table S1). Both of these tumor types are variable for a range of characteristics including size, morphology, color, and vascularization. For example, HCC B from *CAG-A3B* #1 and HCC from *CAG-A3B* #2 from independent animals show differential morphology, colorization, and vasculature.

**Figure 4.**
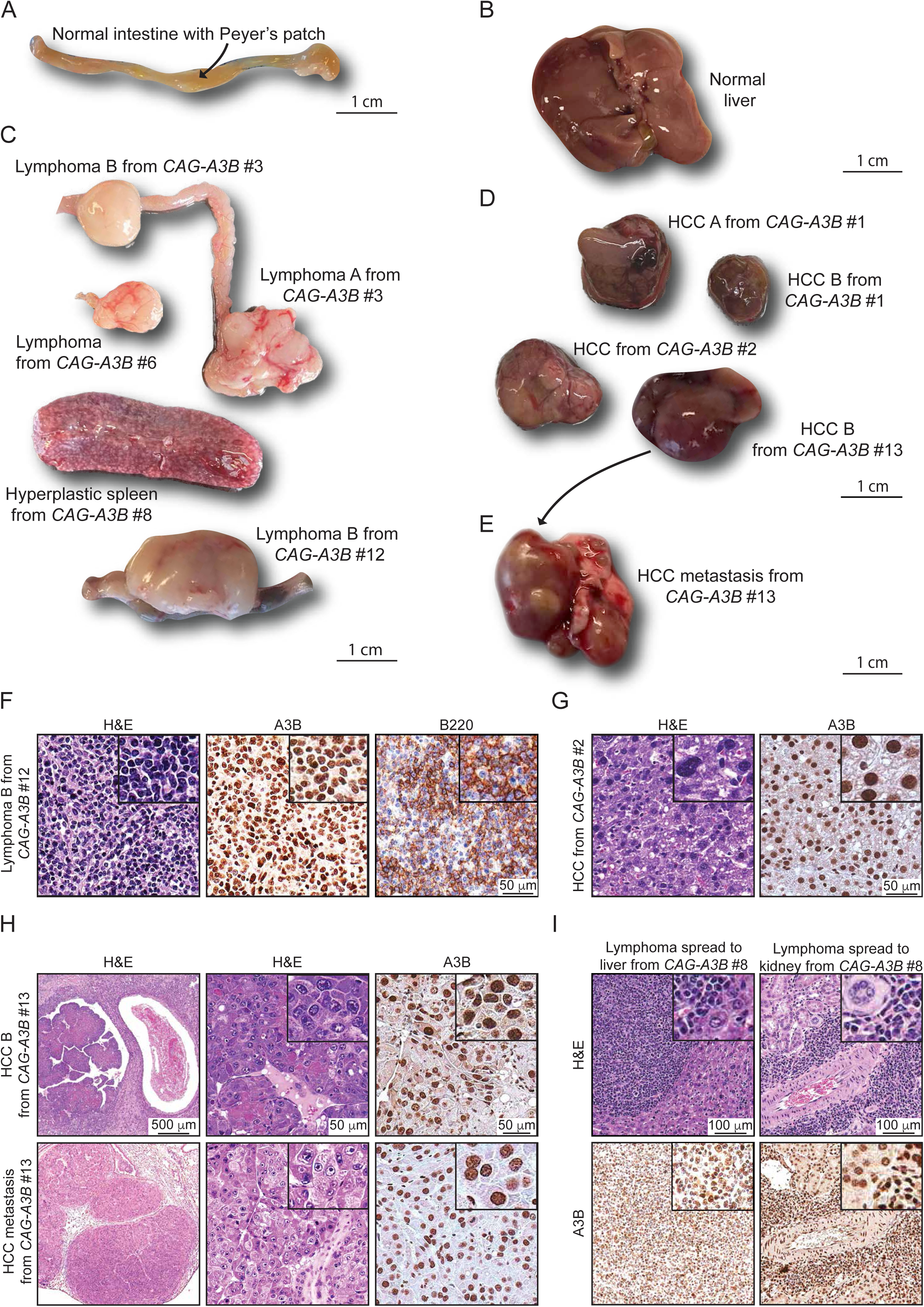
Heterogeneity and evidence for metastasis in tumors from *CAG-A3B* animals. (**A-B**) Representative normal intestine with Peyer’s patch (arrow) and normal liver tissues, respectively, from *CAG-A3B* mice. (**C-D**) Macroscopic pictures of a heterogeneous assortment lymphomas and hepatocellular carcinomas, respectively, from *CAG-A3B* mice. (**E**) Representative image of a primary hepatocellular carcinoma that metastasized to the lung (HCC B from *CAG-A3B* #13 in panel D). (**F**) H&E, anti-A3B, and anti-B220 IHC of lymphoma B from *CAG-A3B* #12. Inset boxes show the same tumors at 4x additional magnification. (**G**) H&E and anti-A3B IHC of HCC from *CAG-A3B* #2. Inset boxes show the same tumors at 4x additional magnification. (**H**) H&E and anti-A3B IHC staining of a primary hepatocellular carcinoma (top) and its metastatic dissemination to the lung (bottom) from *CAG-A3B* #13. Inset boxes show the same tumors at 4x additional magnification. (**I**) H&E and anti-A3B IHC staining of a diffuse large B-cell lymphoma in the liver (left) and kidney (right). Inset boxes show the same tumors at 4x additional magnification. See also **Figures S5** and **S6**.

We next characterized tumors at the cellular level by H&E and IHC for select diagnostic markers. First, all tumors showed diffuse, strong, nuclear-only A3B staining in the entirety of the lesional cells (**Figures 4F-I** and **S5A-B**). Most lymphomas appear to be comprised of a uniform proliferation of atypical lymphoid cells with round or ovoid hyperchromatic nuclei showing marked nuclear pleomorphism, increased number of mitotic figures, and scant eosinophilic cytoplasm (**Figures 4F** and **S5A**). A fraction of lesions also appear macroscopically as enlarged spleens with features suggestive of splenic lymphoid hyperplasia and variable increases in the number and size of follicular structures (**Figures 4C** and **S5B**). Second, staining with the diagnostic B-cell marker B220 indicates that the *CAG-A3B* mice are developing predominantly B-cell lymphomas, either *de novo* or from preceding lymphoid hyperplasias (**Figures 4F** and **S5A-B**). This inference is supported by the clonality of antibody gene contigs derived from RNAseq data, indicating that tumorigenesis occurs after V(D)J recombination in the B-cell lineage (**Figure S5C**). However, several B-cell lymphomas are accompanied by abnormally large, mostly non-clonal, T-cell populations as determined by CD3 staining, *Thy-1* mRNA levels, and diverse TCR junctions, which may be the result of strong anti-tumor T-cell responses and/or inflammation in the tumor microenvironment (**Figure S5A**-**D**).^25^ Notably, *CAG-A3B* HCCs also manifest higher levels of the DNA damage marker γ-H2AX in comparison to adjacent normal liver tissue, which is consistent with ongoing chromosomal DNA deamination by A3B (**Figure S5E-F**). In contrast to near-uniform A3B staining, only a subset of cells is positive for γ-H2AX staining suggesting an involvement of other factors such as cell cycle stage.

Importantly, a subset of *CAG-A3B* animals also show evidence of distant organ metastasis, or disseminated lymphoproliferative malignancy, with one case of HCC metastasizing to the lung, one case of disseminated lymphoma involving the Peyer’s patches and intestinal muscosa, two cases of lymphoma with diffuse lymph node dissemination to multiple lymph nodes, and one case of lymphoma spreading to multiple lymph nodes, the liver, and the kidney (*e.g.*, **Figures 4C**, **4E**, **4H-I** and **S6**). In several instances, both the primary and the metastatic lesions are located adjacent to blood vessels (**Figure 4H-I**). In the case of liver-to-lung metastasis, the metastatic tumor shows indistinguishable histopathologic features from the primary HCC and similarly uniform and strong A3B positivity (**Figure 4H**). In agreement with this, all disseminated lymphoproliferative lesions also show strong A3B nuclear-only immunostaining (**Figures 4I** and **S6**), which differs from our prior studies in which human A3A protein expression is selected against and disappears in early stages of HCC development.^15^ No metastases were observed in the WT mice over the same time frame. These observations combine to suggest that A3B influences both early- and late-stage tumor development in an ongoing manner.

### *CAG-A3B* tumors exhibit APOBEC signature mutations

Whole genome sequencing (WGS) of tumors from *CAG-A3B* and WT animals, in comparison to matched tail DNA, enables somatic mutation landscapes to be compared and underlying mutational processes to be deduced. First, tumors from both *CAG-A3B* and WT animals exhibit variable numbers of all types of single base substitution mutations (*CAG-A3B*: n=29 tumors, SBS range 1111-67414, mean=12893, median=3304; WT: n=9 tumors, SBS range 1538-8203, mean=5016, median=6101; **Figure S7A**). Second, visual comparison of the distributions of single base substitution mutations reveals a larger proportion of C-to-T mutations in most tumors from *CAG-A3B* mice in comparison to tumors from WT animals (representative examples in **Figure 5A** and all additional profiles in **Figures S8 and S9**). These mutations occur predominantly in TCA, TCC, and TCT trinucleotide motifs, consistent with the established biochemical and structural preferences of A3B in which the target cytosine is most frequently preceded by thymine.^10, 26–29^ Indeed, this TC-focused C-to-T mutation bias is known as SBS2, and the calculated percentage of SBS2 mutations in *CAG-A3B* tumors associates positively with overall APOBEC mutation signature enrichment score (red data points in **Figure S7B**). In contrast, neither SBS2 nor significant APOBEC mutation signature enrichment is observed in tumors from WT animals, which is expected because short read WGS predominantly identifies clonal (or near-clonal) somatic mutations (black data points in **Figure S7B**).

**Figure 5:**
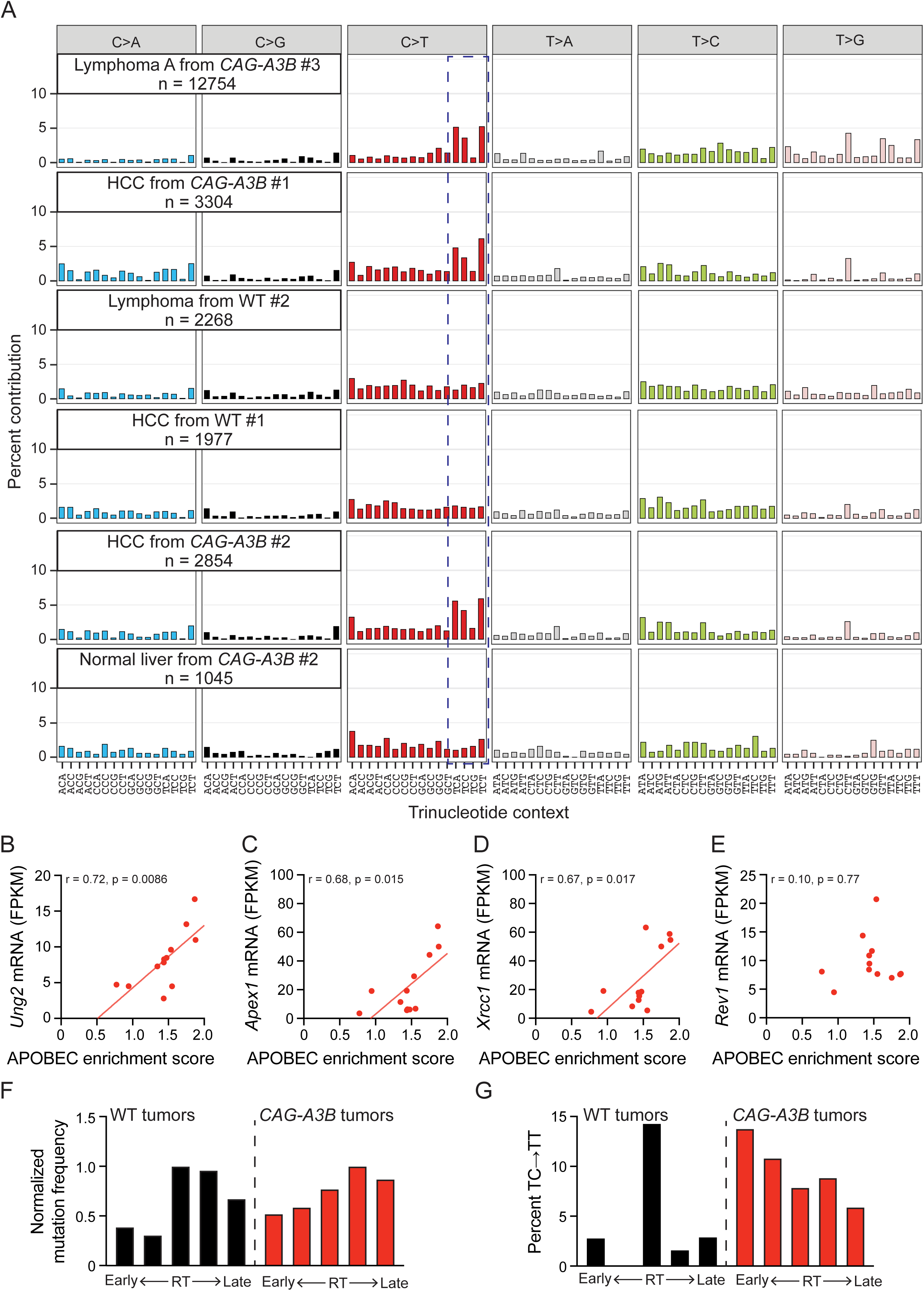
*CAG-A3B* tumors exhibit APOBEC signature mutations. **(A)** Representative SBS mutation profiles for the indicated tumors from WT or *CAG-A3B* animals (mutation numbers shown). The dashed box highlights APOBEC preferred TC motifs characteristic of SBS2. (**B-E**) Scatterplots of APOBEC enrichment score from *CAG-A3B* lymphomas (n=12) compared to the mRNA levels of *Ung2*, *Apex1*, *Xrcc1*, and *Rev1*, respectively, from the same tumors (Pearson correlation coefficients and corresponding p-values indicated). (**F**) Bar plots showing the proportion of mutations in WT and *CAG-A3B* tumors according to early-to late-replicating regions (mutation numbers normalized to the largest quintile in each group). (**G**) Bar plots showing the percent of T<u>C</u>-to-T<u>T</u> mutations as a percent of all mutations in each quintile in panel F. See also **Figures S7**, **S8**, **S9**, **S10**, and **S11**.

Curiously, C-to-G transversion mutations in the same TC-focused trinucleotide motifs (SBS13) are not apparent above background levels in tumors from *CAG-A3B* animals (representative examples in **Figure 5A** and additional profiles in **Figures S8 and S9**). SBS2 and SBS13 are thought to be alternative mutational outcomes of APOBEC3-catalyzed C-to-U deamination events, with the latter signature attributable to uracil excision by uracil DNA glycosylase (Ung2) followed by C-insertion opposite the newly created abasic site by the DNA polymerase Rev1. An analysis of the mRNA levels of uracil excision repair factors in *CAG-A3B* lymphomas indicates positive associations between APOBEC signature enrichment and *uracil DNA glycosylase 2* (*Ung2*), *AP endonuclease 1* (*Apex1*), and *X-ray repair cross-complementing protein 1* (*Xrcc1*), but not with *Rev1* (**Figure 5B-E**). Conversely, WT lymphomas lack an association between these transcripts and APOBEC enrichment score and, in the case of *Apex1*, may even exhibit a negative association (**Figure S10A-D**). Thus, the absence of SBS13 in *CAG-A3B* tumors may be explained by elevated rates of error free repair (fewer persisting abasic sites) and/or insufficient Rev1 levels (more opportunities for DNA polymerases that follow the A-rule to misincorporate dAMP opposite a lesion).

It is further notable that the vast majority of A3B-associated C-to-T mutations are dispersed and not occurring in small or large clusters called *omikli* and *kataegis*, respectively. A *de novo* extraction of the single base substitution mutation signatures in the entire set of murine tumors yields seven distinct signatures including one closely resembling SBS2 (Sig D in **Figure S11**). This analysis also reveals other candidate mutational signatures including Sig E and Sig G, which most closely resemble SBS17 (oxidative damage) and SBS5 (unknown process associated with aging) in humans (**Figure S11**). Additionally, increases in APOBEC signature mutations also associate with higher mutation burdens in tumors from *CAG-A3B* mice but not in tumors from WT mice (**Figure S7C**). The overall distribution of single base substitution mutations also appears to differ between *CAG-A3B* tumors and WT tumors with the former showing a bias toward genomic regions associated with early replicating timing (**Figure 5F**). This mutation bias for early replicating regions becomes even more pronounced when only TC-to-TT SBS2 mutations are analyzed (**Figure 5G**).

### Hypermutated *CAG-A3B* tumors also exhibit higher frequencies of a range of structural variations

Given prior reports,^30, 31^ we were also interested in determining whether A3B causes structural variation. One can easily imagine how a subset of A3B-catalyzed deamination events may become abasic sites, ssDNA nicks, and dsDNA breaks and be processed into a wide variety of different non-SBS mutagenic outcomes. We first quantified small-scale events ranging from single nucleotide insertion and deletion mutations (indels) to larger-scale indels <200 bp (**Figure 6A**). Interestingly, single T/A indels and single C/G indels are visibly elevated in *CAG-A3B* tumors in comparison to control tumors from WT animals (**Figure 6A-E**), analogous to a recent report with human A3A overexpression in a chicken cell line.^32^ In comparison, no significant differences are seen with 2, 3, and 4 bp indels or with 2, 3, and 4 deletions with microhomology, which might reflect the relatively small number of events in each of these categories (**Figure 6A** and **6F-I**). Accordingly, indels in the larger 5+ category (5 bp to 200 bp) are elevated in *CAG-A3B* tumors in comparison to control tumors from WT animals (**Figure 6A**, **6J-K**). As anticipated from these mostly positive results, the total sum of all of these indel events in *CAG-A3B* tumors associates positively with the APOBEC signature enrichment scores consistent with these events sharing a mechanistic origin (**Figures 6L** **and S7D**).

**Figure 6.**
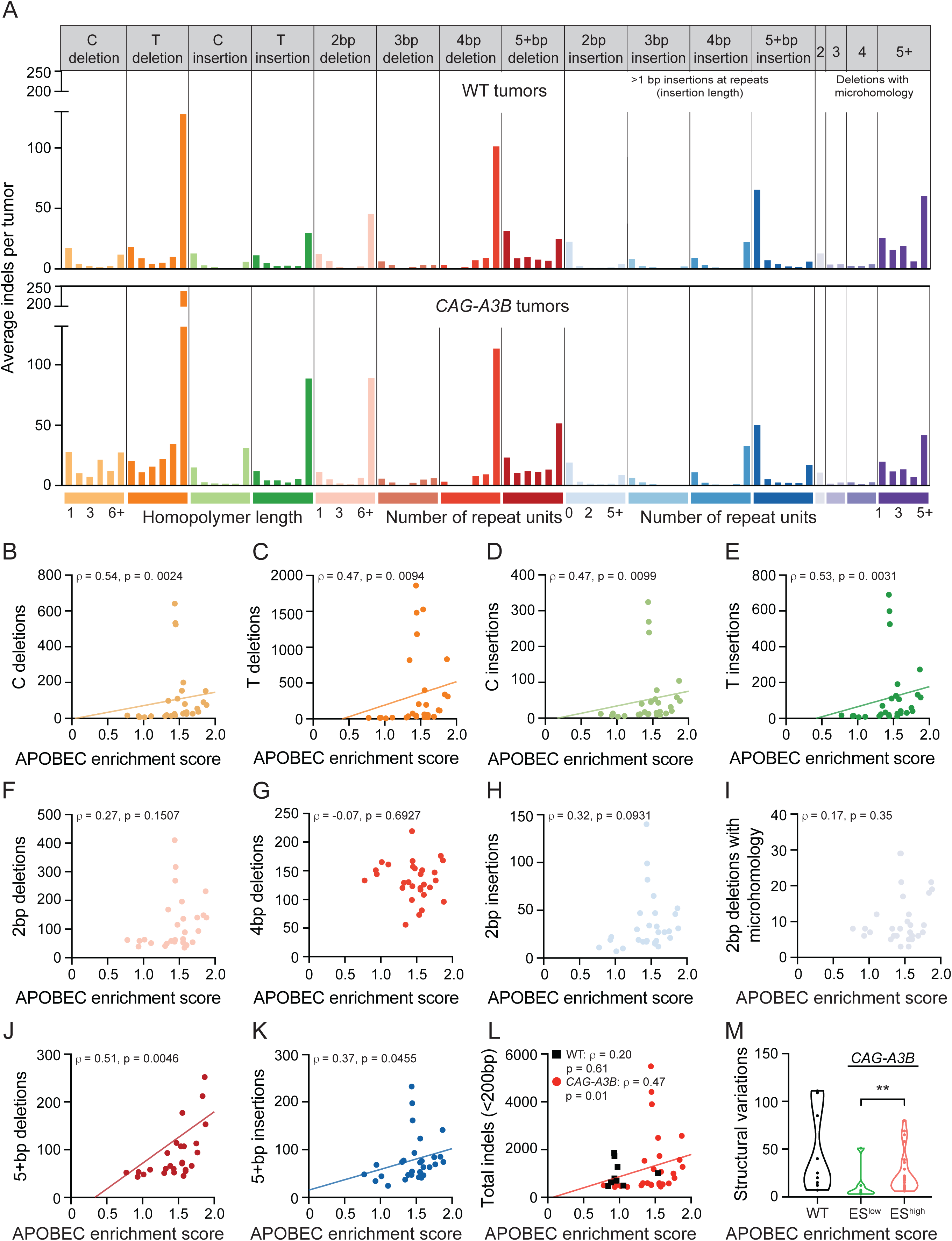
Hypermutated *CAG-A3B* tumors also exhibit higher frequencies of a range of structural variations. (**A**) Composite spectrum of the average number of small insertion/deletion mutations in tumors from WT (n=9) and *CAG-A3B* (n=29) animals. (**B-K**) Scatterplots showing relationships between APOBEC enrichment scores from *CAG-A3B* tumors and the indicated indel types (Spearman’s rank correlation coefficients and corresponding p-values indicated). (**L**) Scatterplots showing the relationship between APOBEC enrichment scores from WT (black) and *CAG-A3B* (red) tumors and the total number of indels <200bp in each tumor (Spearman’s rank correlation coefficients and corresponding p-values indicated). (**M**) Violin plots of the total number of structural variations in tumors from WT mice in comparison to tumors from *CAG-A3B* animals with low or high APOBEC enrichment scores (ES; p=0.0087 for ES^high^ vs ES^low^ groups by Mann-Whitney U test). See also **Figures S7**, **S10**, and **S12**.

On an even greater scale, one must also consider larger structural variations including indels >500 bp, inversions, translocations, and more complex events. We therefore quantified these structural variations in tumors from WT mice and in tumors from CAG-A3B mice with low and high APOBEC enrichment scores (ES^low^ and ES^high^, respectively). Interestingly, a statistically higher level of structural variation is evident in tumors with ES^high^ in comparison to those with ES^low^ (**Figure 6M**). There is a wide range in the number of structural variations in WT mice, and in some tumors, these are nearly as high as in ES^high^ tumors (**Figure 6M**). However, it should be noted that these tumors originate in animals that are much older than those from the *CAG-A3B* cohort, and also that there is a linear correlation between age of the mouse and structural variations in WT tumors consistent with a different mutational mechanism (**Figure S12A**). Additionally, this is true for all mutations and all indels, making it difficult to compare numbers in tumors from WT mice to those in tumors from *CAG-A3B* mice (**Figure S12B-C**). Chromosomal copy number variations were not measured due to the near-isogenicity of the animals enrolled in our studies.

## Discussion

These studies are the first to demonstrate that human A3B drives tumor formation *in vivo* by accelerating rates of primary tumor development as well as by triggering secondary growths (*i.e*., metastases). Specifically, full body expression of CAG promoter-driven levels of human A3B, which approximate those reported in many human tumors, results in accelerated rates of B-cell lymphomagenesis and hepatocellular carcinogenesis as well as a smaller number of other tumor types. Nearly all A3B-expressing tumors also exhibit significant macroscopic heterogeneity as well as elevated levels of C-to-T mutations in TC dinucleotide motifs (SBS2) consistent with the established biochemical activity of this ssDNA deaminase. Importantly, A3B-driven tumors also show positive associations between APOBEC mutation signature enrichment and multiple types of indel mutations. These observations are consistent with a model in which some C-to-U DNA deamination events lead to signature C-to-T mutations and others are processed by uracil base excision repair enzymes into ssDNA breaks that can be converted into indels.^9, 26, 33^ A significantly elevated level of structural variation is also apparent in tumors with high APOBEC signature enrichment in comparison to those with low enrichment. Altogether these results support a model in which tumor development can be both initiated and fueled by a wide range of genetic changes inflicted by A3B (**Figure 7**).

**Figure 7.**
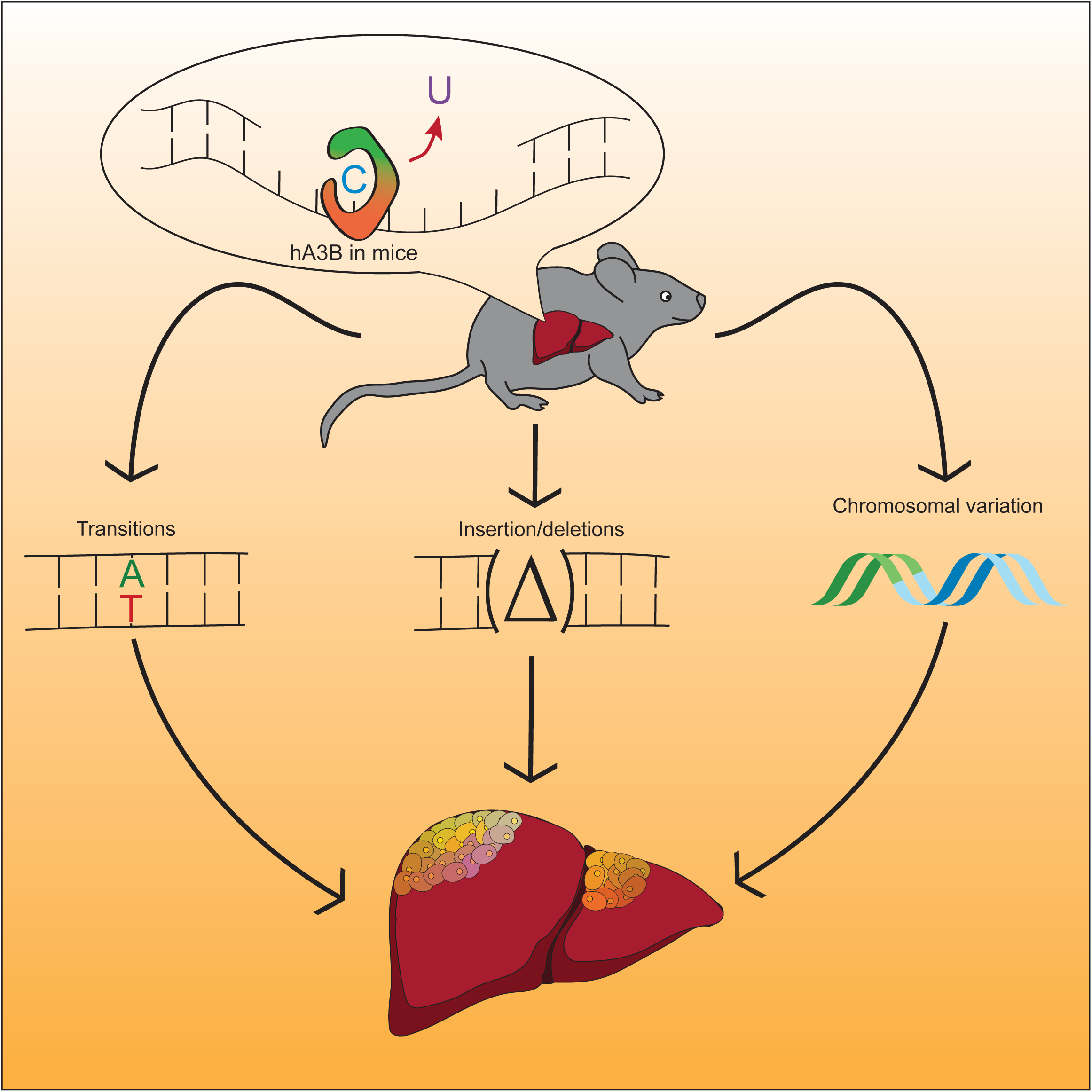
Working model for mutation and carcinogenesis by human A3B. Human A3B catalyzes ssDNA C-to-U deamination events that lead to signature SBS events, as well as small-scale insertion/deletion mutations and larger-scale structural variations. These different mutational events combine to initiate primary tumor development, cause the observed heterogeneity, and fuel additional tumor evolution including metastases.

These studies also led to two major unexpected results. First, only C-to-T mutations characteristic of SBS2 are evident in A3B-expressing murine tumors, and not C-to-G mutations characteristic of SBS13. These two mutation signatures frequently coincide in human tumors but can occur separately as reported for B-cell lymphoma cell lines, urothelial carcinomas with micropapillary histology, Apc^Min^ colorectal tumors in A3A transgenic mice, and for yeast and human cells defective in uracil DNA glycosylase or the translesion DNA polymerase Rev1^11, 15, 34–37^. A possible molecular mechanism is suggested by low *Rev1* expression levels in A3B-expressing tumors here and a lack of an association between these mRNA levels and APOBEC mutation signature enrichment. If Rev1 is not present to insert C opposite abasic sites (downstream of A3B deamination of C and Ung2 excision of the resulting U), then a DNA polymerase that follows the A-insertion rule is likely to substitute and contribute to the observed C-to-T transition mutation bias.

Second, A3B-expressing males (but not females) are completely sterile. Our studies indicate that testes and sperm are morphologically normal, and that the defect manifests post-fertilization between the 2- and 4-cell stage of development. This result is consistent with the DNA damage and genetic instability reported above and additional studies will be needed to delineate the precise defect(s). This result contrasts with most male-specific infertilities, which manifest as underdeveloped testes^37–39^. It is additionally curious that males are affected specifically and that whole-body A3B-expressing females have thus far been fertile for >20 generations.

Since the first implication of A3B mutagenesis in breast cancer in 2013(ref.26), there has been an urgent need to develop a robust mouse model for mechanistic and preclinical studies. Our first attempt resulted in transgene inactivation, most likely by A3B selecting against itself and promoting its own inactivation.^15^ Our second used the endogenous *Rosa26* promoter to drive human *A3B* expression.^16, 41^ This leads to modest A3B expression levels in most murine tissues, normal fertility, no overt cancer phenotypes, and subtle effects in lung-specific cancer models (without APOBEC signature mutations) in the presence of drug selection.^16^ Our third, reported here, leads to higher, human tumor-like levels of A3B in most murine tissues. These “just right” or “Goldilocks” levels of human A3B catalyze genomic instability and accelerate rates of tumorigenesis and, for as-yet-unknown reasons, only trigger male and not female infertility. On the higher end of the expression spectrum, Dox-induced expression of human A3B in mice leads to multiple pathological phenotypes and death of all animals within 10-12 days.^21^ Thus, because most normal human tissues express very low (or no) A3B,^26, 42–44^ an open and important question is how can tumors evolve to tolerate high A3B expression levels and associated genomic instabilities without dying?

The Cre-inducibility of the CAG-A3B tumorigenesis model described here may be helpful for fine-tuning the tissue-specific expression of this DNA mutating enzyme and modeling additional human cancer types that show high frequencies of APOBEC signature mutations. The Goldilocks expression levels of human A3B enabled by the CAG promoter (after Cre-mediated excision of the transcription stop cassette) are likely to be helpful for studying the formation, evolution, and therapy responsiveness of many different APOBEC signature-high tumor types including those of the bladder, breast, cervix, head/neck, lung, and other tissue types. Moving forward, it is now clear that both human A3A^15^ and A3B (this study) can drive tumor formation, and should be considered “master drivers” because the mutations they inflict are incredibly heterogeneous and, in turn, affect a constellation of processes including all of the classical hallmarks of cancer. Future mechanistic and preclinical studies focused on tumor diagnosis and therapeutic treatment of APOBEC mutation signature-positive tumors must now endeavor to carefully address both of these carcinogenic enzymes.

## ACKNOWLEDGEMENTS

We thank M. Burns and E. Law for early contributions to this project and D. Bacich for thoughtful feedback. We also thank Y. You and Mouse Genetics Laboratory at the University of Minnesota for help with *in vitro* fertilization experiments, C. Guo and the Gene Targeting & Transgenic Facility at the HHMI Janelia Campus for assistance in establishing the *CAG-A3B* mouse line, the University of Minnesota Genomics Center for next-generation sequencing, U. Randen at the Akershus University Hospital and staff at the University of Minnesota Veterinary Diagnostic Laboratory for help with immunohistological analyses. Cancer studies in the Harris lab are supported by NCI P01-CA234228 and a CPRIT Established Investigator Recruitment Award. Murine model development initiated with support from a Jimmy V Foundation Translational Grant, a Norwegian Centennial Chairs Seed Grant (grant ID2016/12240), a Randy Shaver Cancer Research and Community Fund seed grant, and a Department of Defense CDMRP Idea Award (BC121347 to R.S.H.). C.D. was supported briefly by NIGMS T32-GM140936. Salary support for R.L.-K. was provided by NHLBI T32-HL007062 and subsequently by NCI F32-CA232458. R.L.-K. is an awardee of the Weizmann Institute of Science National Postdoctoral Award Program for Advancing Women in Science. HN is supported by the Research Council of Norway (grant no. 229633), Norwegian Cancer Society (grants no. 2017/2715;223314-2022), and the Southeast Regional Health Authority (project no. 274901). RSH is an Investigator of the Howard Hughes Medical Institute, a CPRIT Scholar, and the Ewing Halsell President’s Council Distinguished Chair at University of Texas Health San Antonio.

## AUTHOR CONTRIBUTIONS

C.D., R.L.-K., and R.S.H. conceptualized the overall project. C.D., R.L.-K., L.A., S.C.H., A.A.V., J.P., A.H., and Z.S. conducted *in vivo* mouse experiments. C.D., R.L.-K., P.P.A., A.A.V., J.P., A.H., and Z.S. performed molecular biology experiments. C.D., N.A.T., Y.T.L., and R.I.V. provided statistical analyses. C.D., R.L.-K., N.A.T., R.S., H.N., and R.S.H. contributed to funding. C.D. and R.S.H. drafted the manuscript with all authors contributing to manuscript proofing and revision.

## STAR METHODS

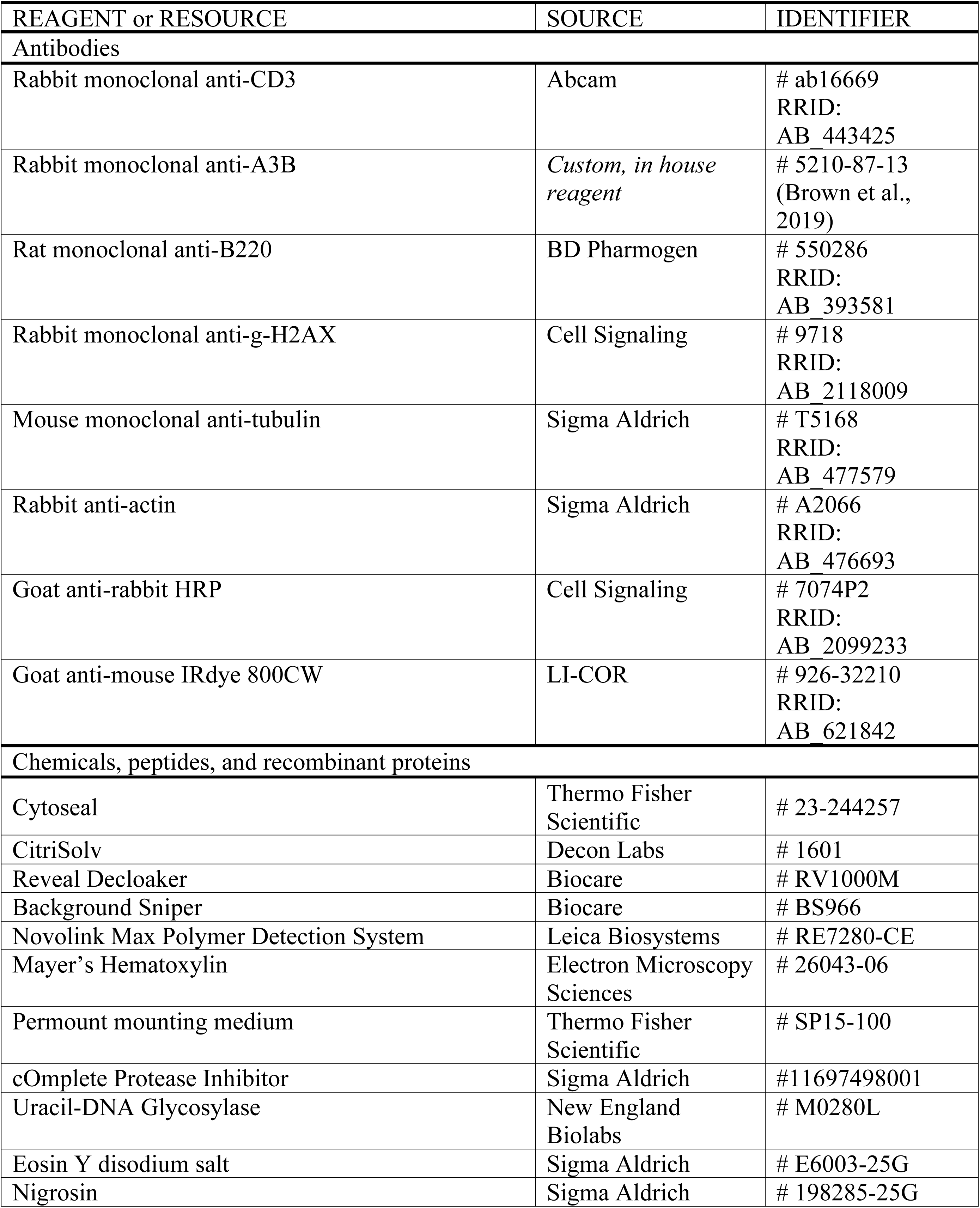
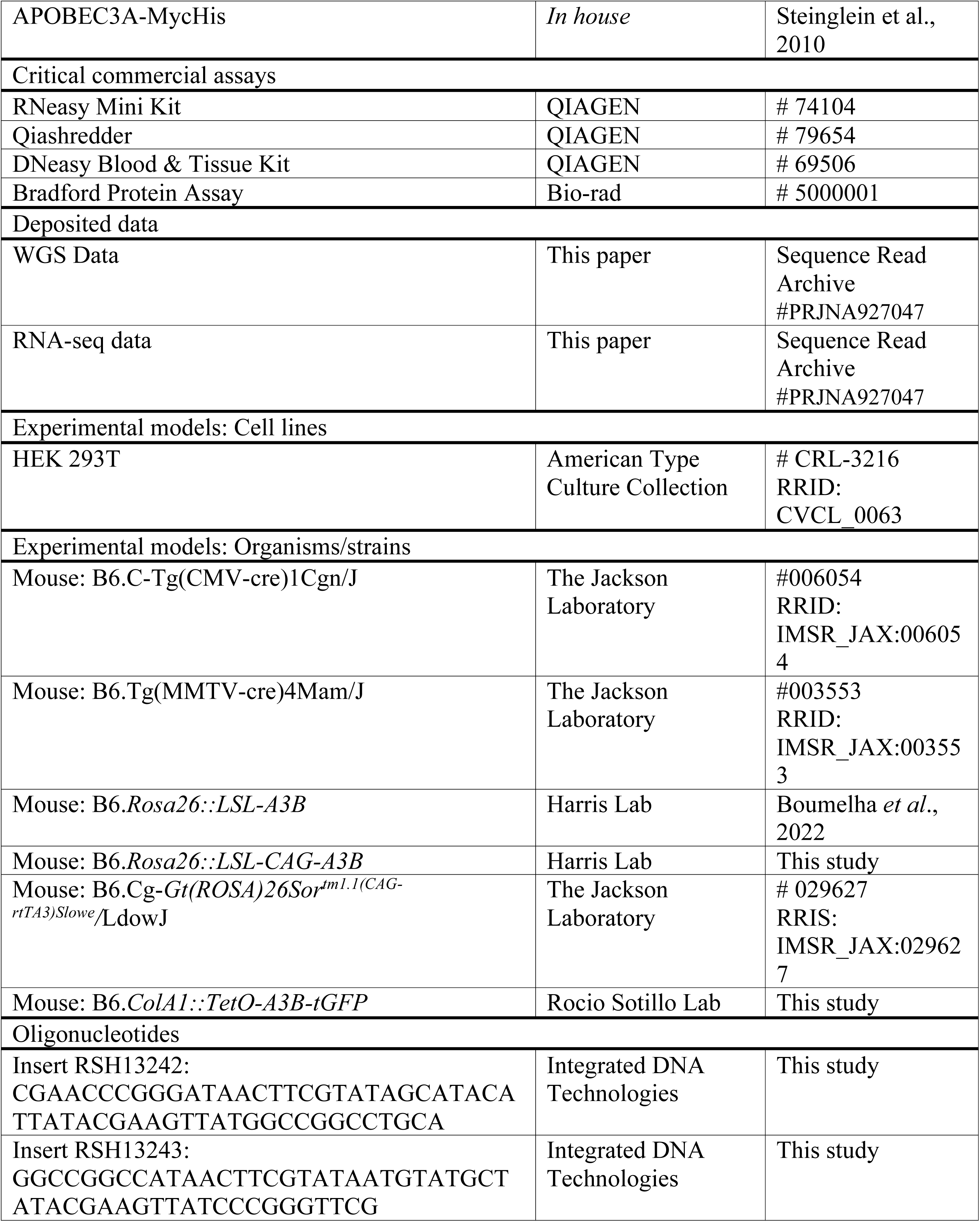
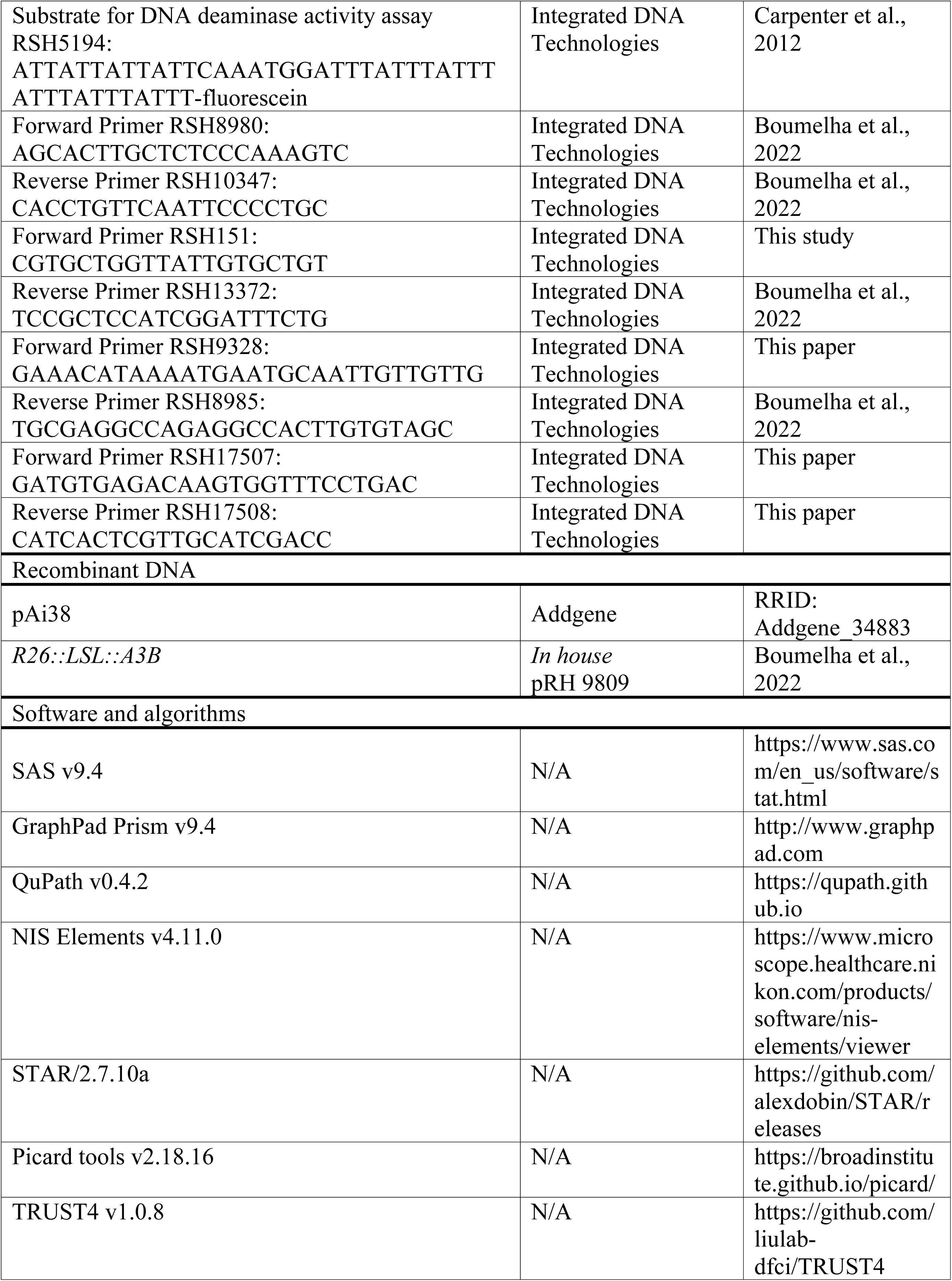
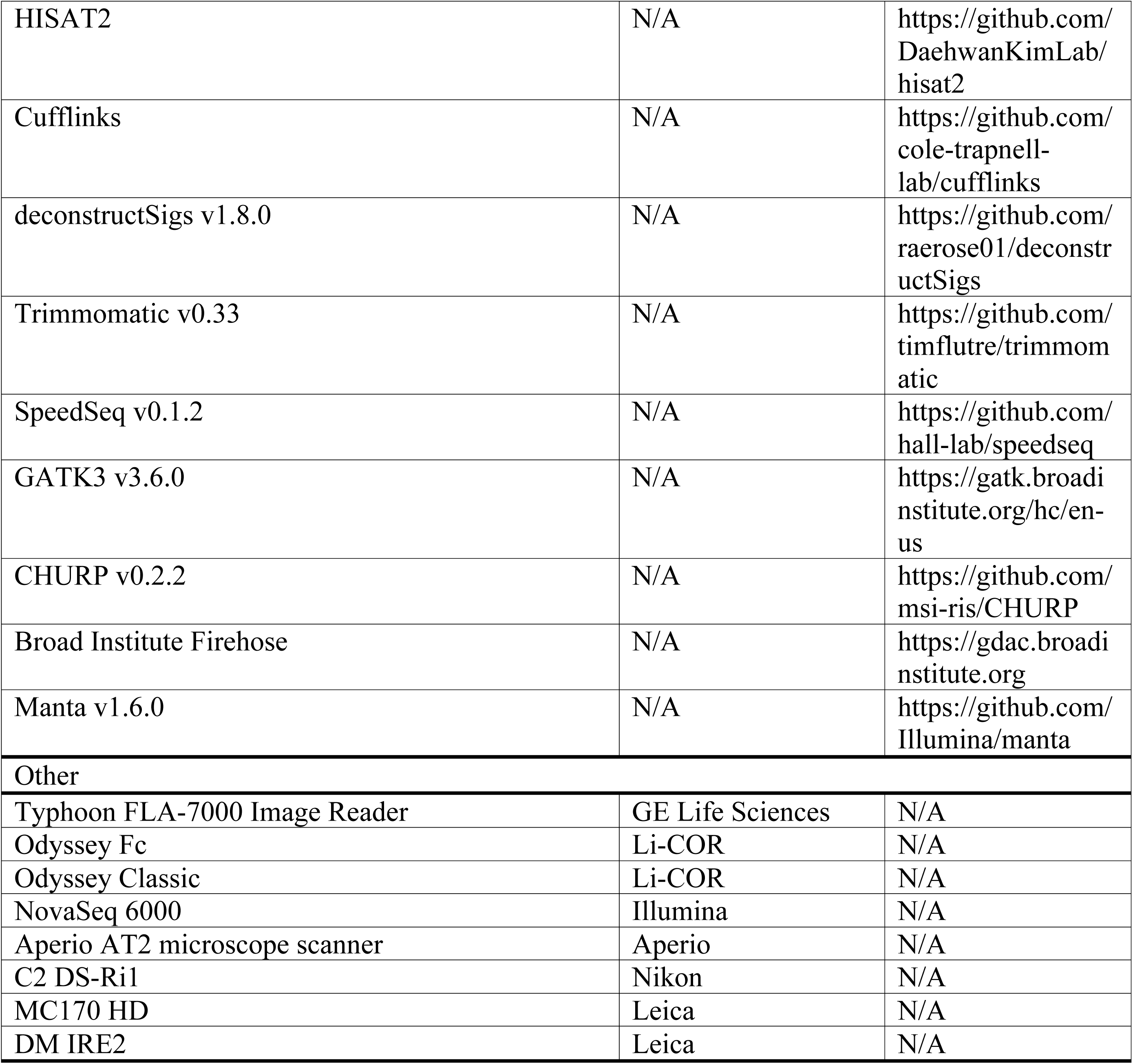
KEY RESOURCES TABLE

## RESOURCE AVAILABILITY

### Lead Contact

Requests for additional data needed to recapitulate results reported in the paper should be directed to the lead contact: Reuben S. Harris.

### Materials availability

Plasmids generated in this study are available through the lead contact. The CAG-A3B animals described here will be deposited at The Jackson Laboratories repository, catalog number pending.

### Data and code availability

Whole genome sequencing and RNA-seq data uploads to the Sequence Read Archive at the National Library of Medicine are in progress, project #PRJNA927047.

## EXPERIMENTAL MODELS AND SUBJECT DETAILS

C57BL/6 mice were used to generate knock-in models at the DKFZ (B6.*ColA1::TetO-A3B-tGFP*) or at the Gene Targeting & Transgenic Facility at the HHMI Janelia Campus (B6.*Rosa26::LSL-CAG-A3B*). Other experimental mice were purchased from The Jackson Laboratories. All mice were housed in specific pathogen-free conditions at 22*°*C under a standard 12 hour light/dark cycle, and handled in agreement with local Animal Care and Ethics committees. All mice were fed standard laboratory chow, with the exception of B6.*ColA1::TetO-A3B-tGFP* mice, which were fed food pellets containing doxycycline (625 p.p.m.; Harlan-Teklad). Mice were housed at the University of Minnesota Twin Cities and University of Texas Health San Antonio animal facilities in specific pathogen-free conditions in accordance with the Institutional Animal Care and Use Committee guidelines (protocol 2201-39748A and 20220024AR, respectively). Mouse experiments performed in DKFZ animal facilities had ethical approval from Baden-Wurttemberg, Germany (license number G-29-19). Murine experiments at the University of Oslo, Institute of Basal Medical Sciences animal facilities were done in minimal disease units and had ethical approval from the Norwegian Food Safety Authority (FOTS ID7569). Both male and female mice were used for all experiments, except for *MMTV-Cre CAG-A3B* mice, which were exclusively female. Mice of both genders developed tumors, and both male and female mice were analyzed using downstream processes including whole-genome sequencing and histopathological analysis. Gender does not affect tumor development in studies described here, although *CAG-A3B* males are infertile. *R26-A3B* mice were reported previously,^16^ and this study provides additional numbers and a more detailed tumor analysis. Additionally, mouse tumor-free survival here only reports animals that were euthanized due to poor body condition.

## METHOD DETAILS

### A3B knock-in

A description of the *R26::LSL-A3B* construct has been published^16^. A *R26::CAG-LSL-A3B* targeting construct was generated using pAi38 (Addgene, 34883) as a backbone, which contains the strong chimeric CAG (cytomegalovirus early enhancer/chicken Β-actin/rabbit β-globin 3’ splice acceptor) promoter and *Rosa26* targeting arms. The plasmid was cut and a hybridized *loxP* oligo pair was ligated (RSH13242 and RSH13243) to this backbone to make an intermediate plasmid. A fragment containing the NEO-stop-loxP-A3Bi elements from the *R26::LSL-A3B* plasmid was cut and ligated into the backbone of the intermediate plasmid to create the final *R26::CAG-LSL-A3B* construct. Cre-dependent expression and activity of A3B was verified by single or co-transfection into HEK 293T cells and subsequent immunoblotting or deaminase activity assay. Final constructs were used for a targeted knock-in into the *Rosa26* locus of C57BL/6 embryonic stem cells at the Gene Targeting & Transgenic Facility at the Howard Hughes Medical Institute Janelia Research Institute.

### Mouse procedures

Female *CMV-Cre* mice (Jax strain 006054)^45^ were crossed with the *Rosa26::LSL-A3B*^16^ and *Rosa26::LSL-CAG-A3B* (these studies) animals. For mammary ductal cell-specific expression of A3B, *Rosa26::LSL-CAG-A3B* mice were crossed with *MMTV-Cre* mice (Jax strain 003551). The resulting pups were genotyped, enrolled, and monitored weekly and aged out until tumors could be observed either visually or by palpation. Genomic DNA was isolated using the Gentra Puregene protocol (Qiagen) on mouse tail biopsies from animals at 21 days of age, with 50 ng of DNA used as a PCR template. A genotyping schematic for *Rosa26::LSL-A3B* and *Rosa26::LSL-CAG-A3B* mice is provided in **Figure S1**. MMTV-Cre mice were genotyped for Cre using RSH17507 and RSH17508. Mice were monitored three times a week for signs of excessive pain or discomfort, or until their tumors reached >1 cm.^3^ All mice were euthanized via CO_2_ asphyxiation, then control tissues and tumors were immediately collected to be fixed in buffered 10% formalin or flash-frozen in liquid nitrogen. Tumors were initially scored based on visual diagnosis, and then subsequently confirmed with histopathological analysis.

### Hematoxylin and eosin (H&E) staining

All tissues were fixed overnight in 10% buffered formalin, and then embedded in paraffin. Fixed tissues were then sectioned into 4 μm slices and mounted onto positively charged adhesive glass slides. After air-drying, they were baked at 60-62*°*C for 20 minutes, washed with xylene for 5 minutes 3 times, soaked in graded alcohols (100% x 2, 95% x 1 and 80% x 1) for three minutes each, and finally rinsed in tap water for 5 minutes. They were then stained with hematoxylin for 5 minutes and rinsed in tap water for 30 seconds, followed by brief submersion in an acid solution and 30-90 seconds in ammonia water. They were then washed with water for 10 minutes, 80% ethanol for 1 minute, counterstained with eosin for 1 minute, dehydrated in graded alcohols followed by xylene as above, and coverslipped with Cytoseal (Thermo Fisher Scientific).

### Immunohistochemistry (IHC)

IHC was performed as described with minor modifications.^17, 46, 47^ Formalin-fixed paraffin-embedded tissues were sectioned into 4-μm-thick slices and mounted on positively charged adhesive slides. They were then baked at 65°C for 20 minutes to deparaffinize and then rehydrated with three consecutive washes in CitriSolv (Decon Labs) for 5 min each followed by graded alcohols as above, followed by a final 5 min wash in running water. Epitope retrieval was performed with the Reveal Decloaker (BioCare Medical) by steaming for 35 minutes with a subsequent 20 minutes off the steamer. Then, slides were washed for 5 min with running water followed by Tris-buffered saline with 0.1% Tween20 (TBST) for 5 minutes. To suppress endogenous peroxidase activity, the slides were soaked in 3% H_2_O_2_ diluted in TBST for 10 minutes, followed by a 5 min rinse in running water. A 15 min soak in Background Sniper (BioCare Medical) was used to block nonspecific binding, with an immediate successive overnight incubation with primary antibody diluted in 10% Sniper in TBST at 4°C. Primary antibodies used for detection were CD3 (Abcam) at a 1:300 dilution, α-A3B/A/G (Brown et al.) at a 1:350 dilution, B220 (BD Pharmogen) at a 1:100 dilution, and γ-H2AX (Cell Signaling) at a 1:200 dilution. Following overnight incubation with primary antibody, samples were washed with TBST for 5 min, then incubated for 30 min with Novolink Polymer (Leica Biosystems). This was developed by application of the Novolink DAB substrate kit (Leica Biosystems) for 5 min, then it was rinsed in water for 10 min, and counterstained for 5 min using Mayer’s hematoxylin solution (Electron Microscopy Sciences). These were dehydrated in graded alcohls and CitriSolv, then cover-slipped with Permount mounting media (Thermo Fisher Scientific). Slides were scanned using an Aperio AT2 microscope slide scanner and analyzed using QuPath.

### DNA deaminase activity assays

Tissues from animals were homogenized and lysed in HED buffer (25 mM HEPES, 5 mM EDTA, 10% glycerol, 1 mM DTT, and 1x cOmplete protease inhibitor [Roche]). Lysates were sonicated for 20 minutes in a water bath sonicator and cleared by centrifugation. Protein concentration was quantified using a Bradford Assay (BioRad) and were normalized to the same amount for the assay. Samples were incubated for 1 hour at 37°C with the HED buffer solution supplemented with 100 μg/ml RNase A, 0.1 U of uracil DNA glycosylase (NEB), 100 μM RSH 5194 and 1x UDG buffer (NEB)^48^. Sodium hydroxide was added to make a 100 mM concentration solution, and incubated at 98°C for 10 minutes, followed by the addition of 1x formamide buffer (80% formamide, 90 mM Tris, 90 mM Boric acid, 2 mM EDTA) and a subsequent 98°C incubation for 10 minutes. Recombinant A3A-MycHis was expressed and purified as described previously to provide a positive control for deamination activity.^49^ Samples were run on a 15% TBE-Urea gel and imaged using a Typhoon 7000 FLA biomolecular imager (GE Healthcare Life Sciences).

### Immunoblots

Tissue lysates were homogenized, lysed, and quantified as above, and then treated with an equal amount of SDS-PAGE loading buffer (62.5 mM Tris-Cl, pH 6.8, 20% glycerol, 7.5% SDS, 5% 2-mercaptoethanol, and 250 mM DTT) and denatured by heating 95°C. Proteins were then separated using an SDS-PAGE gel and transferred to a polyvinylidene Immobilon-FL membrane. Membranes were washed in PBS, then soaked in 5% milk + PBST to block nonspecific binding. The membranes were incubated in a primary rabbit α-human A3A/B/G antibody 1:1,000 (Brown et al.) and mouse α-tubulin 1:10,000 (Sigma Aldrich), or rabbit α-actin 1:5,000 (Sigma Aldrich) at 4°C overnight. Membranes were then washed in PBST six times for 5 minutes each, then incubated for one hour with secondary α-rabbit HRP (Cell Signaling Technology) and goat α-mouse 800 (LI-COR) at 1:10,000 dilutions supplemented with 0.02% SDS. These membranes were then washed 5 times in PBST and one time in PBS for 5 minutes each, then imaged using an Odyssey Classic scanner and Odyssey Fc imager (LI-COR)

### DNA and RNA extraction and sequencing

Genomic DNA was prepared for sequencing from frozen tissues using the DNeasy Blood and Tissue Kit (Qiagen), and RNA was extracted from frozen tissues using the RNeasy Mini kit (Qiagen). In both cases, tissues were homogenized using Qiashredder columns (Qiagen). For DNA sequencing, libraries were sequenced 2 x 150 paired end using a NovaSeq 6000 instrument (Illumina) to get 30x coverage on the genome. Similarly for RNA sequencing, libraries were sequenced 2 x 150 paired end on a NovaSeq 6000 to get 20 million reads per sample. Raw sequence reads were aligned to the mm10 reference mouse genome. All WGS and RNAseq reactions were done at the University of Minnesota Genomics Center.

### Whole genome sequencing analysis

Whole genome sequencing reads were trimmed to remove low quality reads and adapter sequences using Trimmomatic v0.33.^50^ Trimmed reads were aligned to the mouse genome (mm10) using SpeedSeq.^51^ PCR duplicates were removed using Picard (version 2.18.16). Reads were locally realigned around Indels using GATK3 (version 3.6.0) tools. Single base substitutions and small Indels were called relative to the matched normal tissues using Mutect2. SBSs that passed the internal GATK3 filter with minimum 3 reads supporting each variant, minimum 10 total reads at each variant site and a variant allele frequency over 0.05 were used for downstream analysis. Somatic structural variations were detected using Manta following the somatic structural variation described by Manta using sorted and indexed tumor and matched normal bam files.^52^

### RNA sequencing analysis

RNA-seq reads were aligned to mouse genome mm10 using STAR/2.7.1a with basic two pass mode for realigning splice junctions enabled. Picard tools (version 2.18.16) were then used to mark duplicate reads, split CIGAR reads with Ns at the splice junctions. Immune repertoire of lymphomas (V(D)J enrichment) were reconstructed using TRUST4 following its suggested pipeline.^53^ RNA-seq expression levels were calculated using HISAT2^54^ and Cufflinks.^55^ To compare A3B transcript expression levels across tissues, CHURP was used to map and quantify the expression in mouse tissues,^56^ while gene expression profiles from TCGA human tumor tissues were downloaded from the Broad Institute Firehose.

### Somatic mutation analysis

APOBEC enrichment scores were calculated in the R statistical language (version 4.1.2) using variant calls from the sequencing data. First, the data were organized by 1) filtering for single-base substitutions, 2) filtering for C:G base pairs in the reference sequence, 3) removing mutations derived from the mitochondrial genome, and 4) removing C-to-A substitutions, which are not relevant to APOBEC mutagenesis. Next, a 41-base sequence context, consisting of 20 bases up- and down-stream of the mutated position, was extracted from the mm10 reference genome (*BSgenome.Mmusculus.UCSC.mm10* package; version 1.4.3). Finally, APOBEC enrichment scores were computed using the following formula:

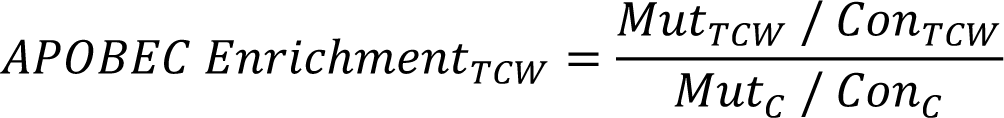

TCW represents the sequence motifs (T<u>C</u>A/T<u>C</u>T) preferred by APOBEC enzymes for cytidine deamination. Mut_TCW_ represents the total number of mutated cytosines in the TCW motif in the 41-base window. Mut_C_ represents the total number of mutated cytosines in the 41-base window. Con_TCW_ and Con_C_ are the total numbers of TCW motifs or cytosines in the 41-base window, respectively. Calculations for the terms above were made for each substitution, and the values were aggregated prior to computing the APOBEC enrichment score for each sample. Statistical significance was calculated using a one-sided Fisher exact test comparing the Mut_TCA_/(Mut_C_ – Mut_TCA_) and the Con_TCW_/(Con_C_ – Con_TCW_) ratios. *P*-values were adjusted using Benjamini-Hochberg correction. The percent contribution of each single-base substitution (SBS) signature (SBS1 to SBS30) was calculated using variant calls from the sequencing data. The *whichSignatures* function in the *deconstructSigs* package (version 1.8.0) in R was applied.

### Male infertility studies

For male sterility experiments, littermate male mice were euthanized, and their cauda epididymis collected. Otherwise, sperm were collected by incubating a lacerated cauda epididymis in a pre-warmed HEPES-0.1% BSA buffer consisting of 130 mM NaCl, 4 mM KCl, 14 mM fructose, 10 mM HEPES, 1.35 mM CaCl_2_, 1 mM MgCl_2_ in a droplet covered by embryo tested neat mineral oil. After incubating the cauda epididymis at 36°C for 30 min to allow the sperm cells to swim out, they were used for the following downstream processes. To take videos of the sperm cells, they remained in the buffer described above while being videoed using a Leica DM IRE2 microscope with the Leica MC170 HD camera. To stain the above sperm cells for a morphological and quantitative viability analysis, eosin-nigrosin staining was performed by using two parts 1% eosin Y (Sigma-Aldrich) and two parts 10% nigrosin (Sigma-Aldrich) well-mixed with one part mouse sperm cells. The resulting mix was then smeared on slides, and a coverslip applied with Cytoseal mounting media. Photographs of these slides were taken using a Nikon C2 DS-Ri1 color camera and analyzed using NIS Elements Viewer. For IVF experiments, a modified version of the Nakagata method was followed,^57^ and developing embryos were quantified and imaged using a Leica DM IRE2 microscope with the Leica MC170 HD camera.

## QUANTIFICATION AND STATISTICAL ANALYSIS

Time to tumor formation was summarized using Kaplan-Meier curves and compared across groups using log-rank tests. Correlative statistical analyses were performed using Pearson correlation coefficient or Spearman’s rank correlation coefficient and were considered significant if the corresponding p-value was <0.05. For statistical analyses to test the outcome between two groups, the median total values were compared by group using Mann-Whitney U test as they have non-normal distributions, while unpaired t-tests were used otherwise. Details for each analysis including test, p-value, and number analyzed can be found in the figure legends or figures themselves. Data were analyzed using SAS 9.4 (Cary, NC) and GraphPad Prism 9.4. P-values <0.05 (or false discovery rate q-values <0.1 for high APOBEC enrichment scores) were considered statistically significant. P-value < 0.05 = *, p-value < 0.01 = **, p-value < 0.001 = ***, p-value < 0.0001 = ****.

