## Supplementary material for "Human APOBEC3B promotes tumor heterogeneity *in vivo* including signature mutations and metastases": Figures S1-S12 and Table S1

[*] Equal secondary contributions.

Supplementary Materials: Figures S1-S12 and Table S1

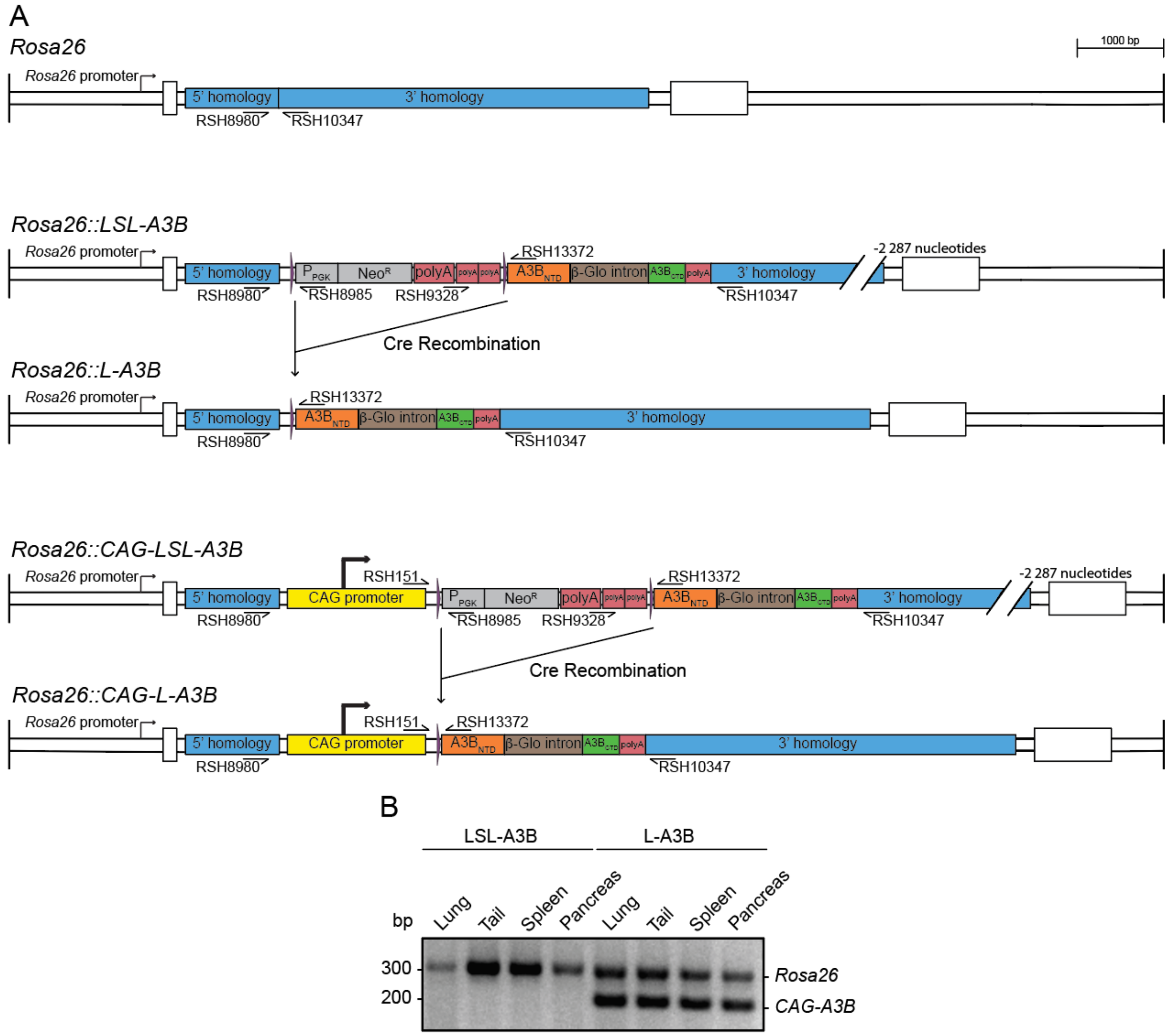


**Figure S1. Construction and validation of mice expressing APOBEC3B, related to Figure 1**

(**A**) To-scale schematics of the WT *Rosa26* locus and, following gene targeting, the fully intact *Rosa26::LSL-A3B* and *Rosa26::CAG-LSL-A3B* minigene alleles. Cre-mediated recombination excises the *loxP*-STOP-*loxP* (LSL) transcriptional stop cassette and allows the *Rosa26* promoter or the *CAG* promoter to drive human *A3B* expression (R26-A3B and CAG-A3B, respectively).

(**B**) PCR genotyping distinguishes the wildtype *Rosa26* locus from each knock-in allele using the indicated primer sets.

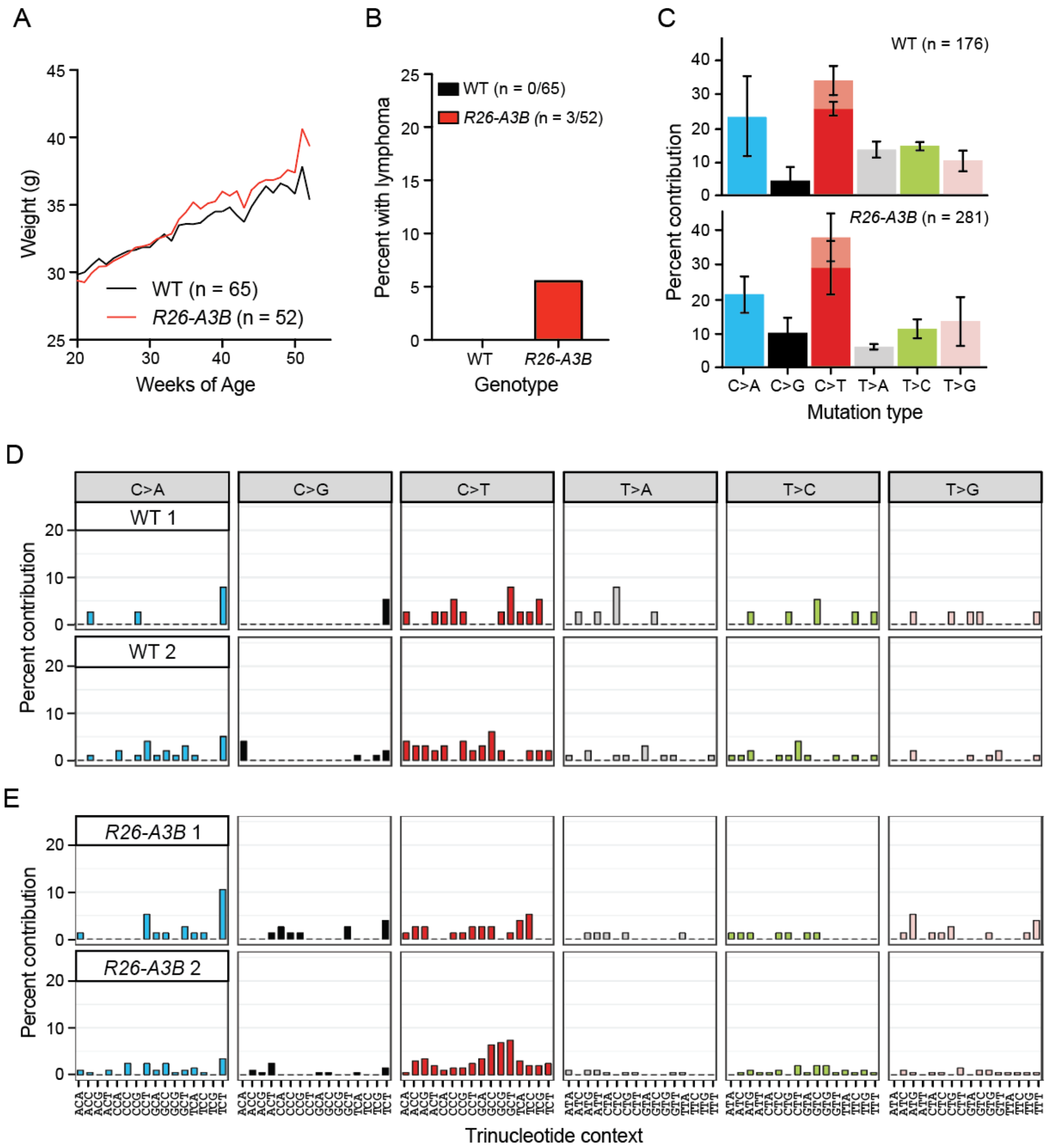


**Figure S2. Low levels of human A3B have limited mutagenic or tumorigenic capacity in mice, related to Figure 3**

(**A**) Average of weights in grams taken weekly from 20-52 weeks of age for each indicated genotype. 65 WT mice and 52 *Rosa26-A3B* mice are represented in this group.

(**B**) Percent of mice between 11 and 23 months that had a lymphoma upon euthanasia and necropsy (p>0.99 by Fisher's exact test).

(**C**) Bar plot showing relative proportion of each type of SBS mutation in spleens from 2 WT (12 months) and 2 *R26-A3B* mice (15 and 18 months), from whole-exome sequencing (WES) at 30x depth. The light red portion of C-to-T mutations represent mutations in CG motifs.

(**D-E**) Trinucleotide mutation profiles of the total SBS mutations in the exomes of WT mice and *CAG-A3B* mice, respectively.

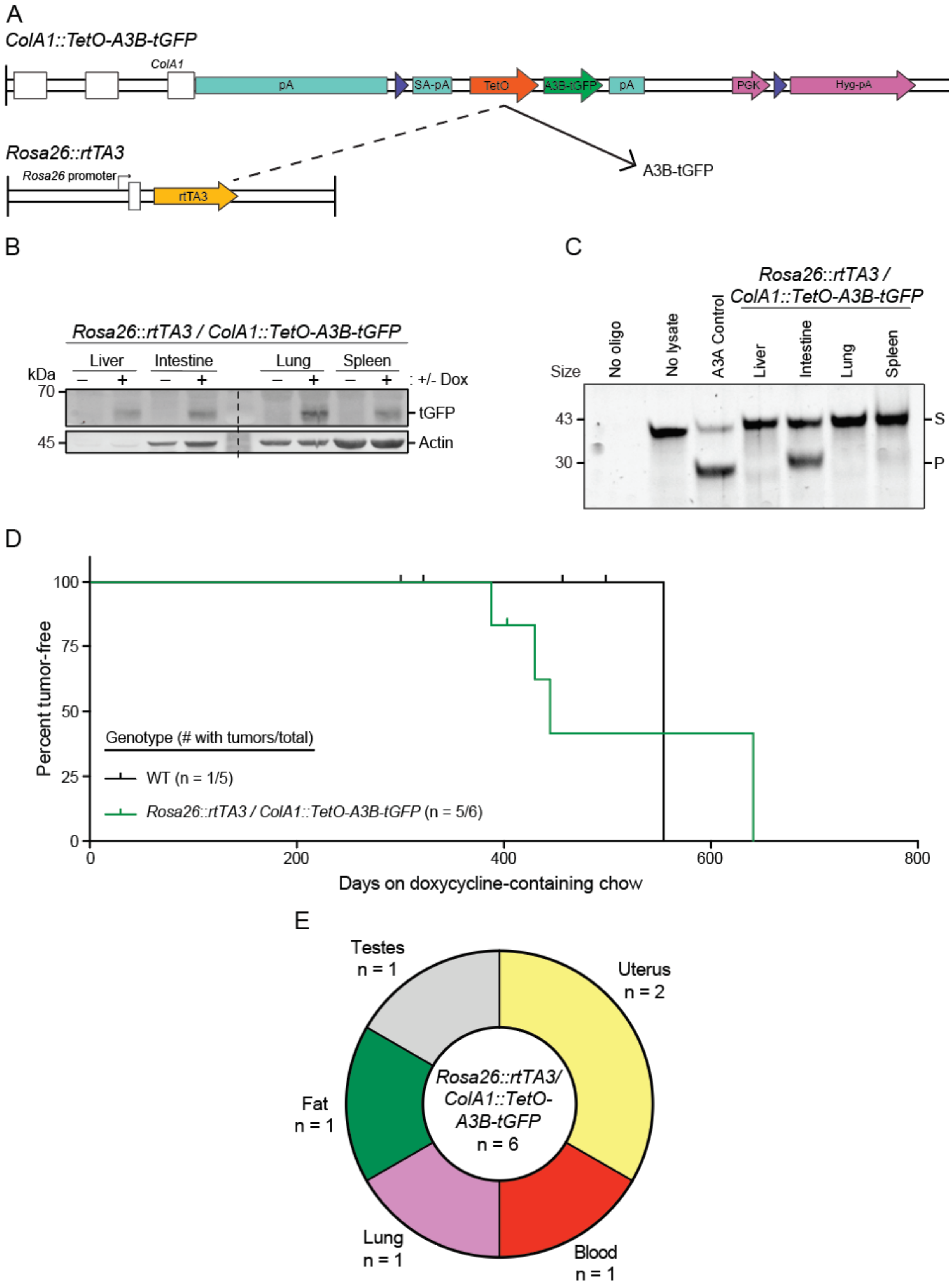
A
ColA1::TetO-A3B-tGFP
ColA1
pA
SA-pA
TetO
A3B-tGFP
pA
PGK
Hyg-pA
Rosa26::rtTA3
Rosa26 promoter
rtTA3
A3B-tGFP
B
Rosa26::rtTA3 / ColA1::TetO-A3B-tGFP
Liver
Intestine
Lung
Spleen
kDa
+/- Dox
70
45
tGFP
Actin
C
Rosa26::rtTA3 / ColA1::TetO-A3B-tGFP
Size
No oligo
No lysate
A3A Control
Liver
Intestine
Lung
Spleen
43
30
S
P
D
100
75
50
25
0
Percent tumor-free
Genotype (# with tumors/total)
WT (n = 1/5)
Rosa26::rtTA3 / ColA1::TetO-A3B-tGFP (n = 5/6)
0
200
400
600
800
Days on doxycycline-containing chow
E
Testes
n = 1
Uterus
n = 2
Fat
n = 1
Rosa26::rtTA3/ ColA1::TetO- A3B-tGFP
n = 6
Lung
n = 1
Blood
n = 1

**Figure S3. A3B expressed from the ColA1 locus is modestly tumorigenic, related to Figure 3**

(**A**) Schematic of the engineered *ColA1::TetO-A3B-tGFP* locus and *Rosa26::rtTA3* locus, which enables expression of A3B-tGFP following doxycycline (Dox) administration.

(**B-C**) Immunoblot and ssDNA deaminase activity of A3B-tGFP expressed in the indicated tissues from *Rosa26::rtTA3/ColA1::TetO-A3B-tGFP* mice fed with (+) or without (-) Dox-containing chow for 15 days. Recombinant A3A is a positive control for activity (S, substrate; P, product), and actin provides a loading control.

(**D**) Kaplan-Meier curves comparing tumor-free survival of WT (n=5) and *Rosa26::rtTA3/ColA1::TetO-A3B-tGFP* (n=6) animals fed with Dox-containing chow for 1-2 years from birth (p=0.37 by log-rank Mantel-Cox test).

(**E**) Pie chart showing the total number of each tumor type found in *Rosa26::rtTA3/ColA1::TetO-A3B-tGFP* mice.

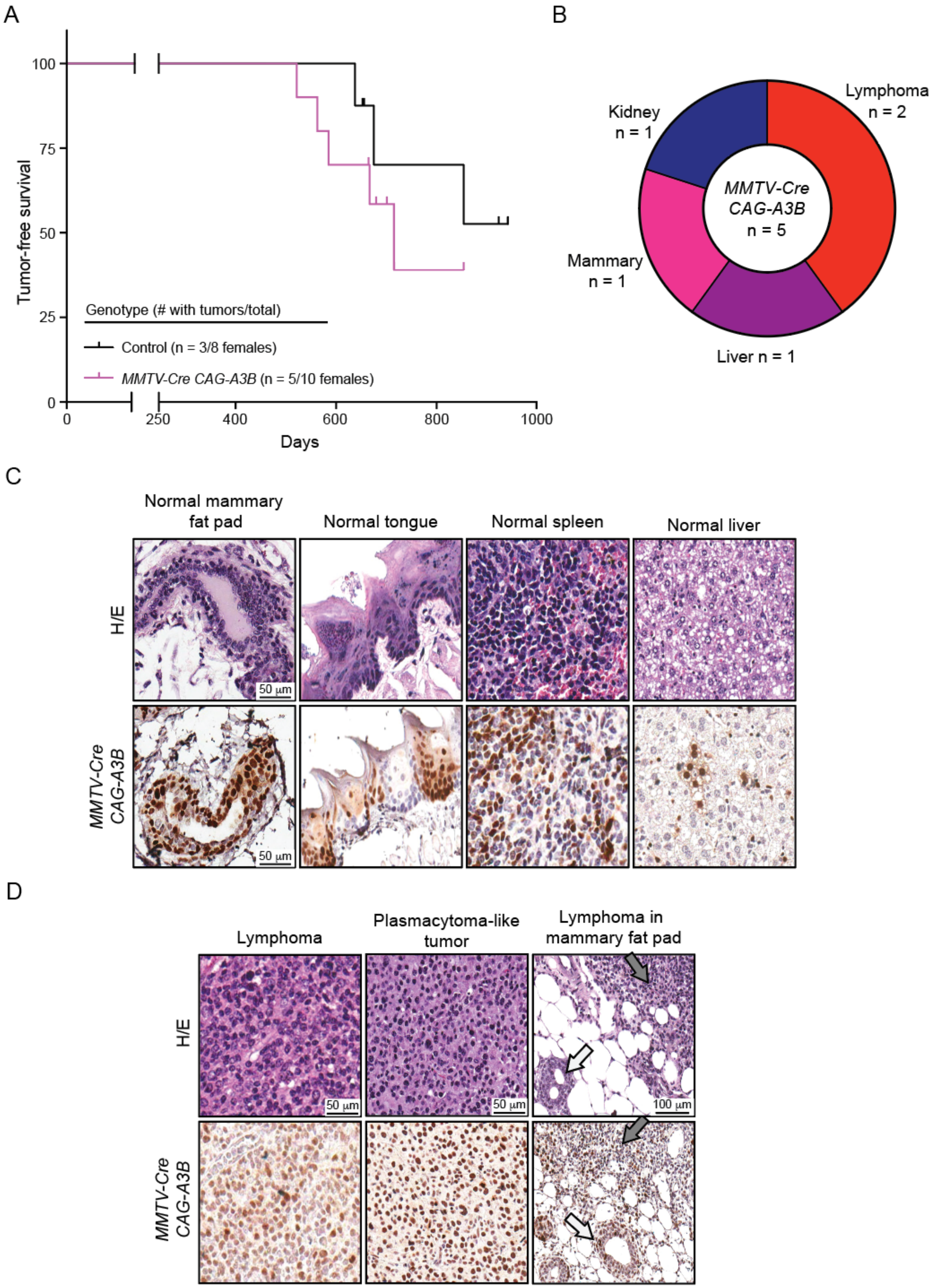
A
100
75
50
25
0
Tumor-free survival
Genotype (# with tumors/total)
Control (n = 3/8 females)
MMTV-Cre CAG-A3B (n = 5/10 females)
0
250
400
600
800
1000
Days
B
Lymphoma
n = 2
Kidney
n = 1
MMTV-Cre
CAG-A3B
n = 5
Mammary
n = 1
Liver n = 1
C
Normal mammary fat pad
Normal tongue
Normal spleen
Normal liver
H/E
MMTV-Cre CAG-A3B
50 μm
50 μm
D
Lymphoma
Plasmacytoma-like tumor
Lymphoma in mammary fat pad
H/E
MMTV-Cre CAG-A3B
50 μm
50 μm
100 μm

**Figure S4. *CAG-A3B* expression mediated by *MMTV-Cre* has limited tumor phenotype, related to Figure 3**

(**A**) Kaplan-Meier curve showing tumor-free survival of *MMTV-Cre* control (n=8 females) and *MMTV-Cre CAG-A3B* (n=10 females) mice (p=0.35 by log-rank Mantel-Cox test).

(**B**) Pie chart showing the total number of primary tumor locations that developed in *MMTV-Cre CAG-A3B* females.

(**C**) H&E and anti-A3B IHC of indicated normal tissues in *MMTV-Cre CAG-A3B* mice. These immunohistological analyses indicate partial *MMTV-Cre* penetrance in multiple tissues beyond the mammary glands.

(**D**) H&E and anti-A3B IHC of representative lymphoproliferative tumors in *MMTV-Cre CAG-A3B* mice. Gray arrows point toward lymphomas, and white arrows to mammary glands.

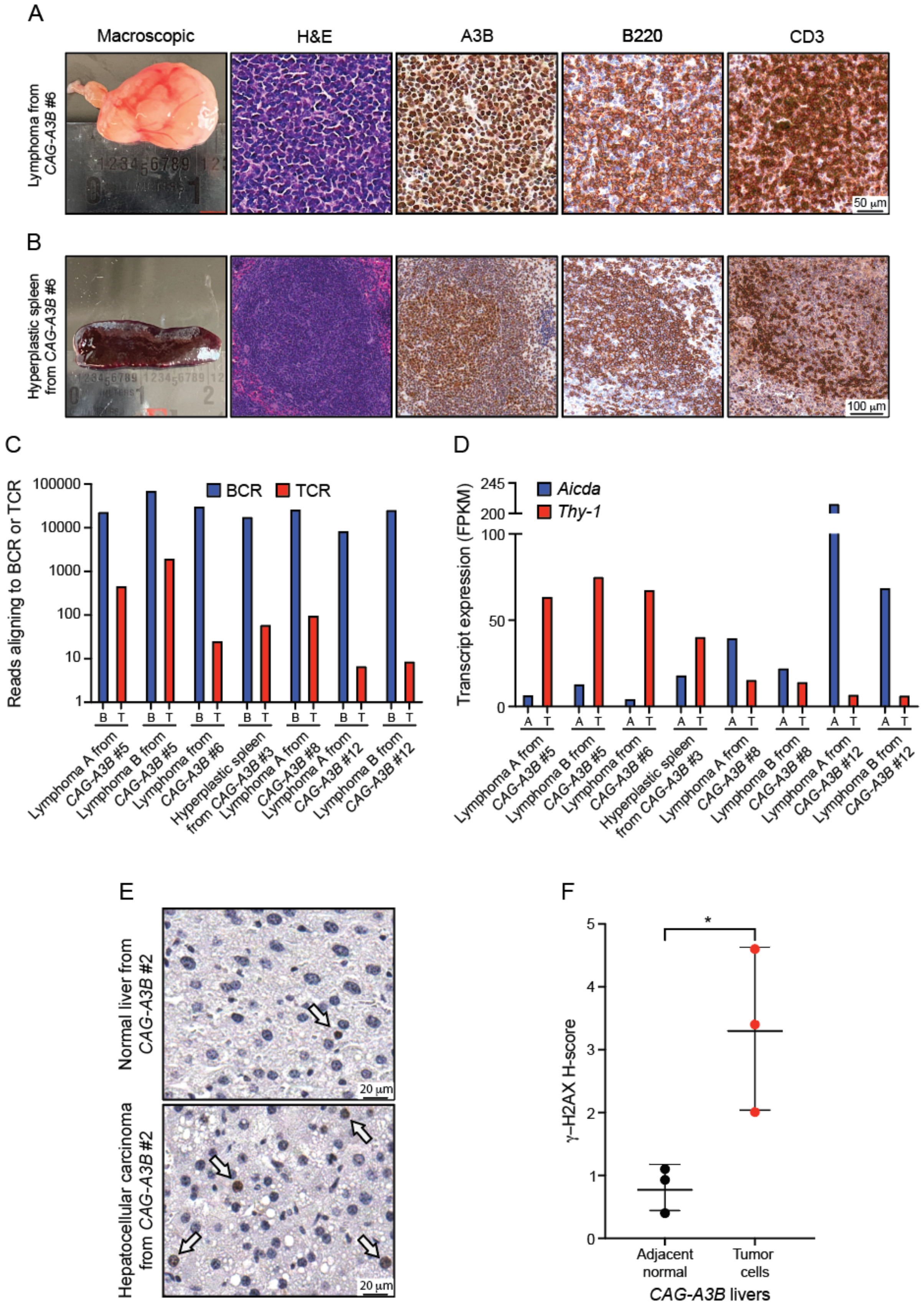
A
Macroscopic
H&E
A3B
B220
CD3
Lymphoma from CAG-A3B #6
50 μm
B
Hyperplastic spleen from CAG-A3B #6
100 μm
C
Reads aligning to BCR or TCR
BCR
TCR
Lymphoma A from CAG-A3B #5
Lymphoma B from CAG-A3B #5
Lymphoma from CAG-A3B #6
Hyperplastic spleen from CAG-A3B #3
Lymphoma A from CAG-A3B #8
Lymphoma A from CAG-A3B #12
Lymphoma B from CAG-A3B #12
D
Transcript expression (FPKM)
Aicda
Thy-1
Lymphoma A from CAG-A3B #5
Lymphoma B from CAG-A3B #5
Lymphoma from CAG-A3B #6
Hyperplastic spleen from CAG-A3B #3
Lymphoma A from CAG-A3B #8
Lymphoma B from CAG-A3B #8
Lymphoma A from CAG-A3B #12
Lymphoma B from CAG-A3B #12
E
Normal liver from CAG-A3B #2
Hepatocellular carcinoma from CAG-A3B #2
20 μm
F
γ-H2AX H-score
Adjacent normal
Tumor cells
CAG-A3B livers

**Figure S5. Phenotypic diversity of *CAG-A3B* tumors, related to Figure 4**

(**A-B**) Macroscopic photograph and photomicrographs of H&E, anti-A3B, anti-B220, and anti-CD3 IHC of the lymphoma from *CAG-A3B* #6 and hyperplastic spleen from *CAG-A3B* #6, respectively. These lesions consist of a mixed B-cell and T-cell population, as shown by high levels of both B220 and CD3 immunostaining. High A3B levels are expressed within the nuclei of most cells. The macroscopic view of the lymphoma is also presented in Figure 4C.

(**C**) A histogram showing the number of *CAG-A3B* lymphoma RNAseq reads that align to recombined B-cell receptor (BCR; B) or T-cell receptor (TCR; T) genes using the TRUST4 algorithm. Each histogram bar represents the largest single contig for each recombined gene. The fact that the BCR rearrangement is clonal and represented by >100-fold more reads demonstrates that most lymphomas have a B cell origin.

(**D**) Transcript expression levels of B-cell specific transcript *Aicda* mRNA and pan T-cell marker *Thy-1* in lymphomas from *CAG-A3B* animals (additional analyses of the same RNAseq data sets as in panel C).

(**E-F**) Representative images and quantification of γ-H2AX immunostaining of HCCs and normal adjacent liver parenchyma from *CAG-A3B* animals, respectively. H-score is determined by the extent of nuclear reactivity in terms of percent cells and is equal to the sum of strong staining = % x 3, medium staining = % x 2, weak staining = % x 1, no staining = % x 0 (n=3 per condition; mean +/- SD shown; p= 0.0314 by unpaired t-test).

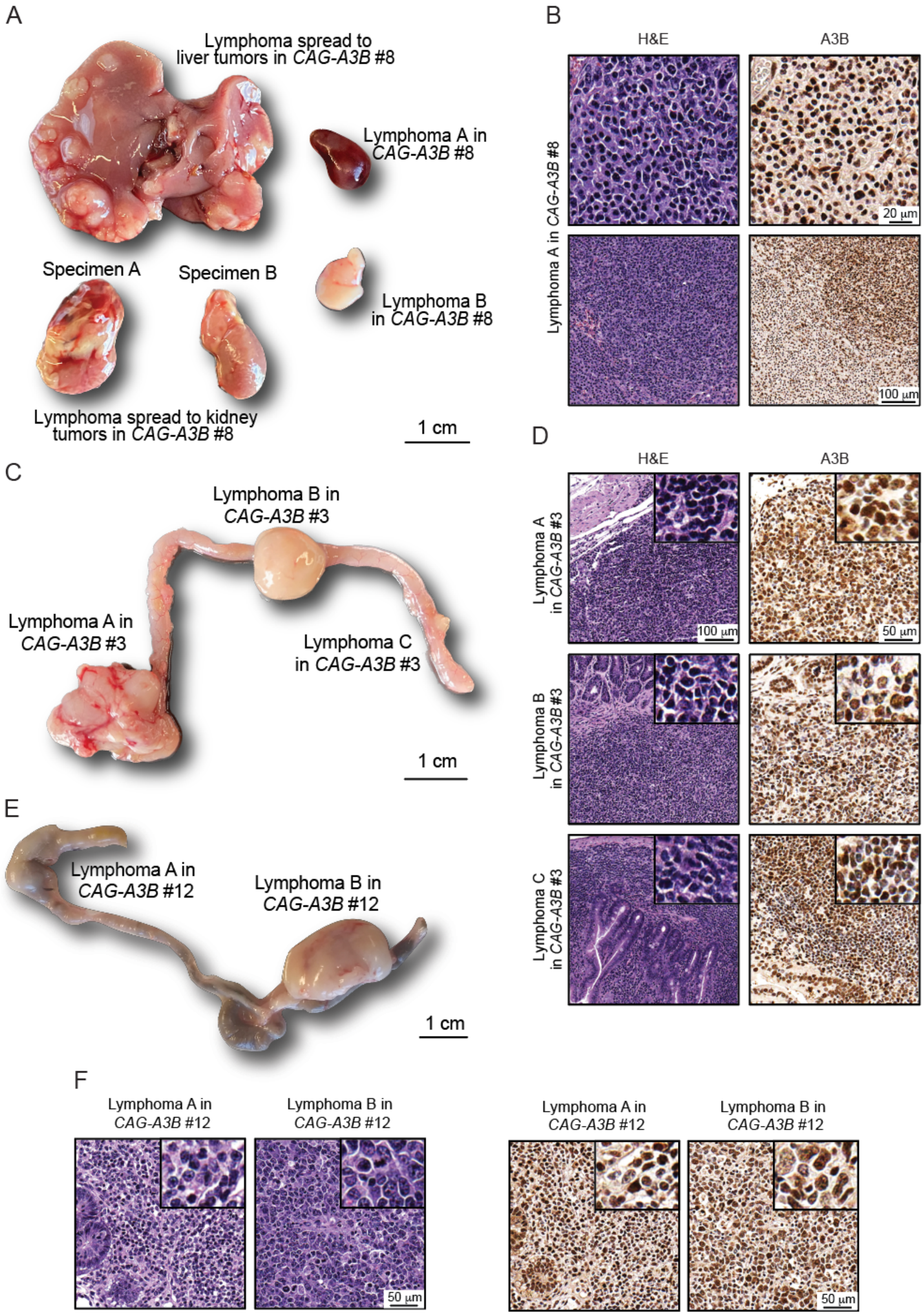
A
Lymphoma spread to liver tumors in CAG-A3B #8
Lymphoma A in CAG-A3B #8
Specimen A
Specimen B
Lymphoma B in CAG-A3B #8
Lymphoma spread to kidney tumors in CAG-A3B #8
1 cm
B
H&E
A3B
Lymphoma A in CAG-A3B #8
20 μm
100 μm
C
Lymphoma B in CAG-A3B #3
Lymphoma A in CAG-A3B #3
Lymphoma C in CAG-A3B #3
1 cm
D
H&E
A3B
Lymphoma A in CAG-A3B #3
100 μm
50 μm
Lymphoma B in CAG-A3B #3
Lymphoma C in CAG-A3B #3
E
Lymphoma A in CAG-A3B #12
Lymphoma B in CAG-A3B #12
1 cm
F
Lymphoma A in CAG-A3B #12
Lymphoma B in CAG-A3B #12
50 μm
Lymphoma A in CAG-A3B #12
Lymphoma B in CAG-A3B #12
50 μm

**Figure S6. Disseminated lymphomas in *CAG-A3B* mice, related to Figure 4**

(**A**) Macroscopic pictures of lymphoma metastasis to the liver, kidney, and multiple lymph nodes in *CAG-A3B* mouse #8.

(**B**) H/E (left) and anti-A3B IHC (right) of lymphoma A in *CAG-A3B* mouse #8. Representative photomicrographs from the lymphoma spread to liver and kidney can be found in Figures 4H-I.

(**C**) Macroscopic picture of a diffuse lymphoma spread to the intestinal mucosa and Peyer's patches in *CAG-A3B* mouse #3.

(**D**) H/E (left) and anti-A3B IHC (right) of lymphomas A-C in *CAG-A3B* mouse #3.

(**E**) Macroscopic picture of a lymphoma involving a mesenteric lymph node and the intestinal mucosa/Peyer's patch in *CAG-A3B* mouse #12.

(**F**) H/E (left) and anti-A3B IHC (right) of lymphomas A and B in *CAG-A3B* mouse #12. All boxed images in panels B, D, and F are representative areas of the corresponding tissues at 4x the magnification. Note that the macroscopic views of the lymphomas in Figure S6C and S6E are also shown in Figure 4C, and shown again here with extended views to more broadly represent all features of the disseminated lymphomas.

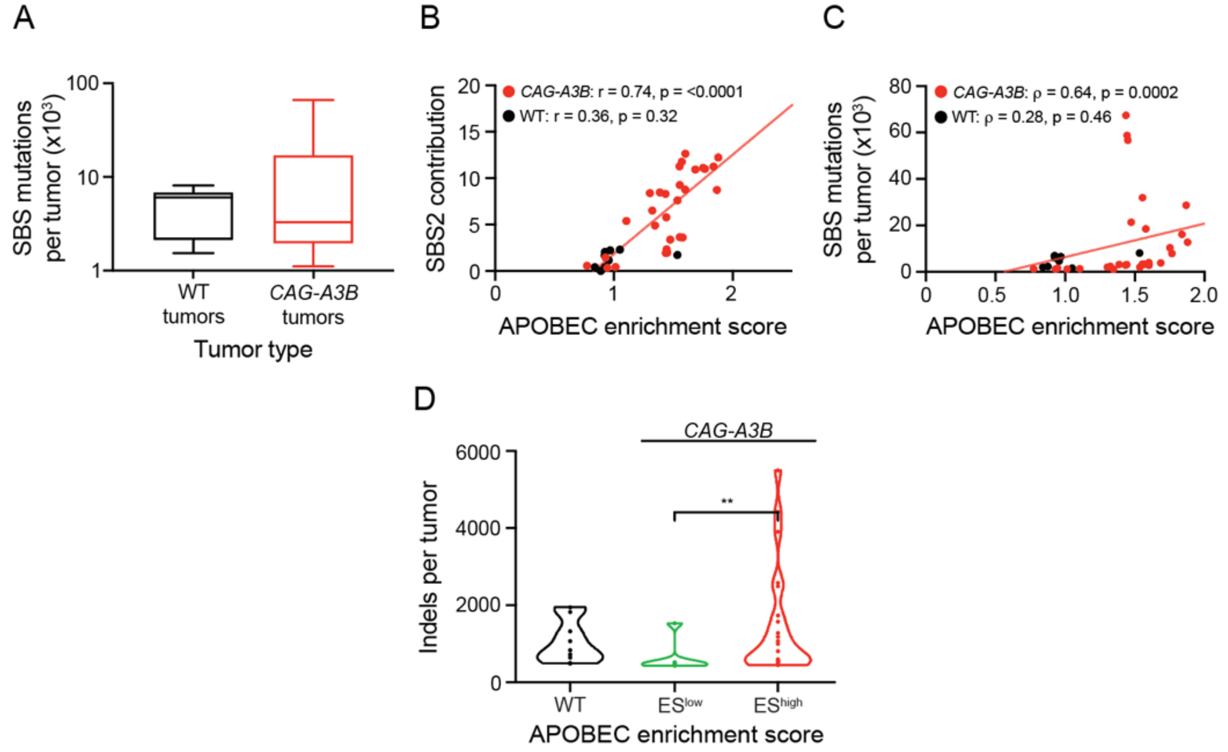


**Figure S7. Genomic mutations in WT and *CAG-A3B* mice, related to Figure 5**

(**A**) Box and whisker plots of the number of SBS mutations in tumors from WT and *CAG-A3B* mice. The middle horizontal line is the median, the outer horizontal lines are the upper and lower quartiles, and the whiskers outside the box represent the maximum and minimum ($p>0.99$ by Mann-Whitney).

(**B-C**) Scatterplots comparing APOBEC mutation signature enrichment scores to the percentages of SBS2 or to the full base substitution loads in tumors from WT and *CAG-A3B* animals, respectively (B, Pearson correlation coefficient, C, Spearman's rank correlation coefficient, and p-values indicated).

(**D**) Violin plots of the total number of indels in tumors from WT mice in comparison to tumors from *CAG-A3B* animals with low or high APOBEC enrichment scores (ES; $p=0.0076$ for $ES^{low}$ vs $ES^{high}$ groups by Mann-Whitney U-test).

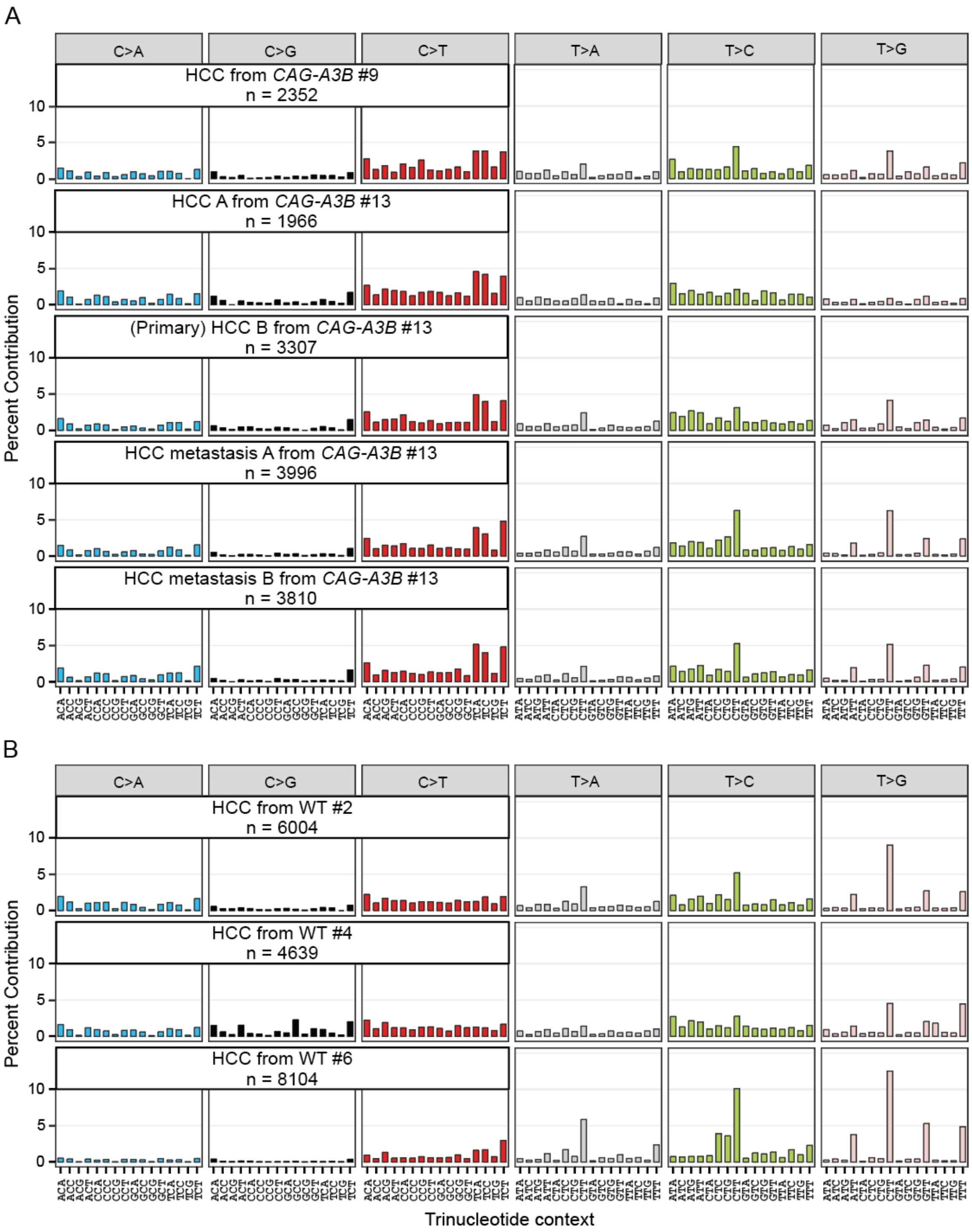


**Figure S8. Mutational spectra of liver tumors, related to Figure 5**

(**A-B**) Trinucleotide mutation profiles of all whole-genome sequenced liver tumors in *CAG-A3B* and WT mice not shown in Figure 5 (n-values = total SBS mutations in each tumor).

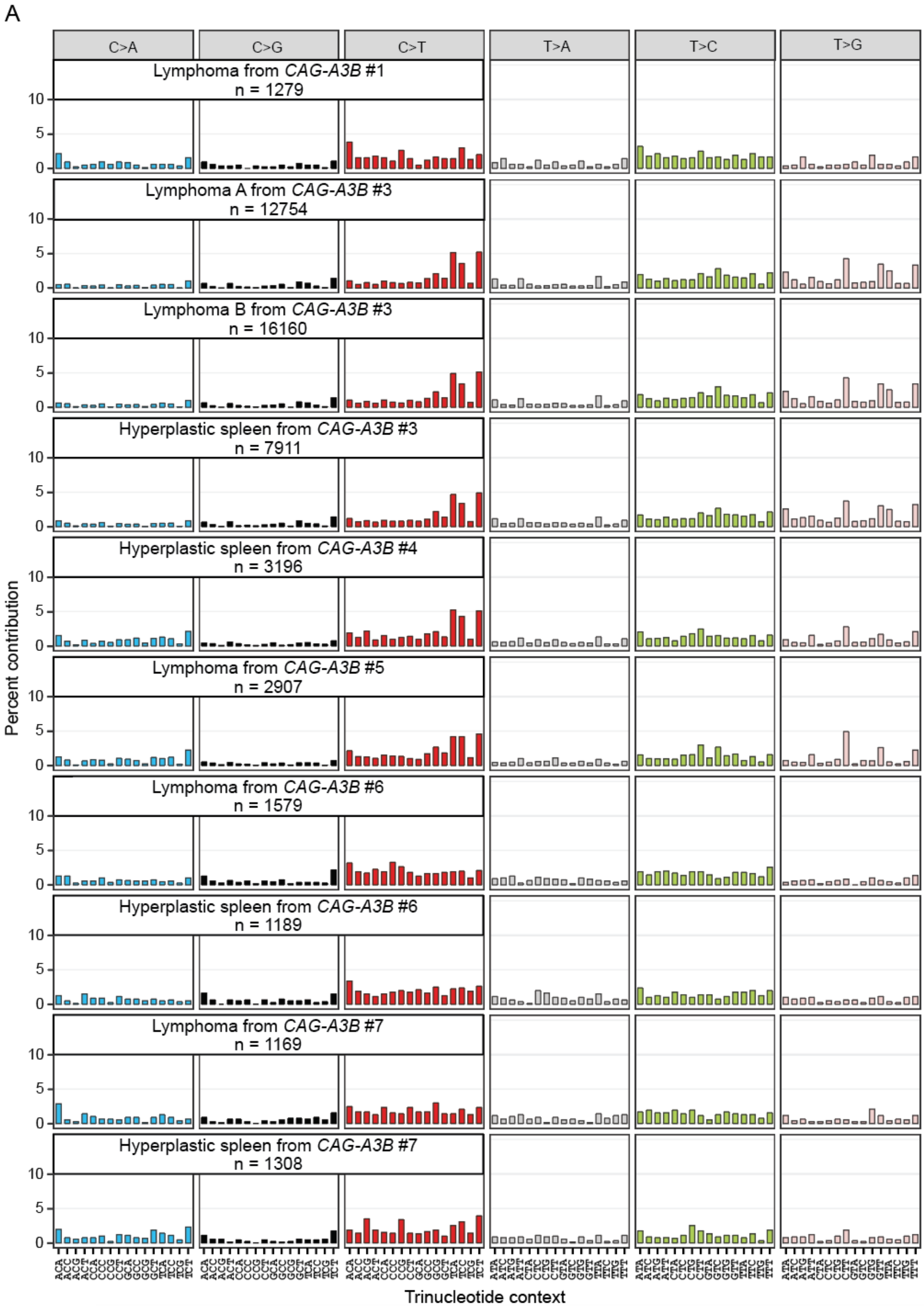
A
C>A
C>G
C>T
T>A
T>C
T>G
Lymphoma from CAG-A3B #1
n = 1279
Lymphoma A from CAG-A3B #3
n = 12754
Lymphoma B from CAG-A3B #3
n = 16160
Hyperplastic spleen from CAG-A3B #3
n = 7911
Hyperplastic spleen from CAG-A3B #4
n = 3196
Lymphoma from CAG-A3B #5
n = 2907
Lymphoma from CAG-A3B #6
n = 1579
Hyperplastic spleen from CAG-A3B #6
n = 1189
Lymphoma from CAG-A3B #7
n = 1169
Hyperplastic spleen from CAG-A3B #7
n = 1308
Percent contribution
Trinucleotide context

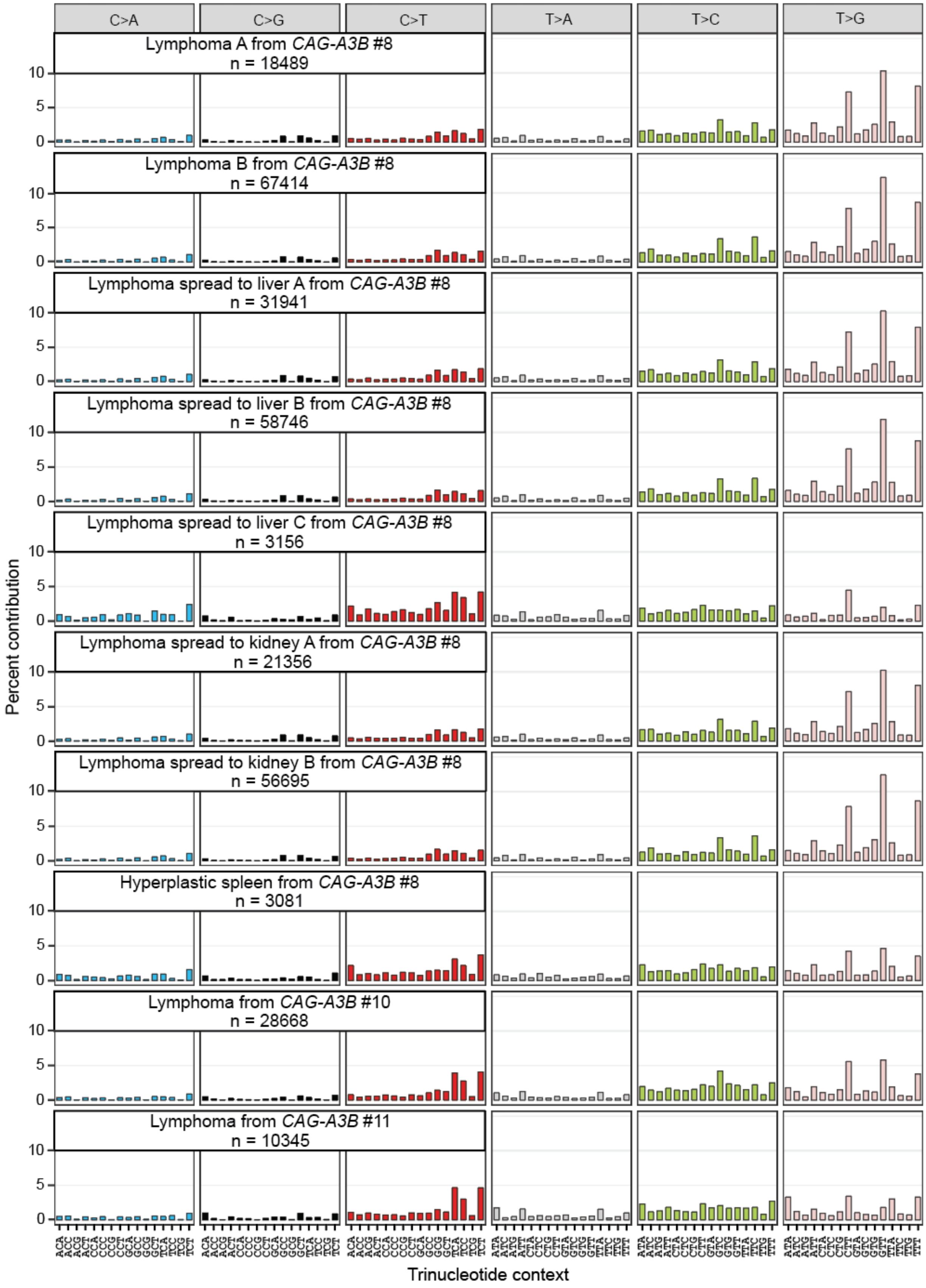
C>A
C>G
C>T
T>A
T>C
T>G
Lymphoma A from *CAG-A3B* #8
n = 18489
Lymphoma B from *CAG-A3B* #8
n = 67414
Lymphoma spread to liver A from *CAG-A3B* #8
n = 31941
Lymphoma spread to liver B from *CAG-A3B* #8
n = 58746
Lymphoma spread to liver C from *CAG-A3B* #8
n = 3156
Lymphoma spread to kidney A from *CAG-A3B* #8
n = 21356
Lymphoma spread to kidney B from *CAG-A3B* #8
n = 56695
Hyperplastic spleen from *CAG-A3B* #8
n = 3081
Lymphoma from *CAG-A3B* #10
n = 28668
Lymphoma from *CAG-A3B* #11
n = 10345
Percent contribution
0
5
10
Trinucleotide context

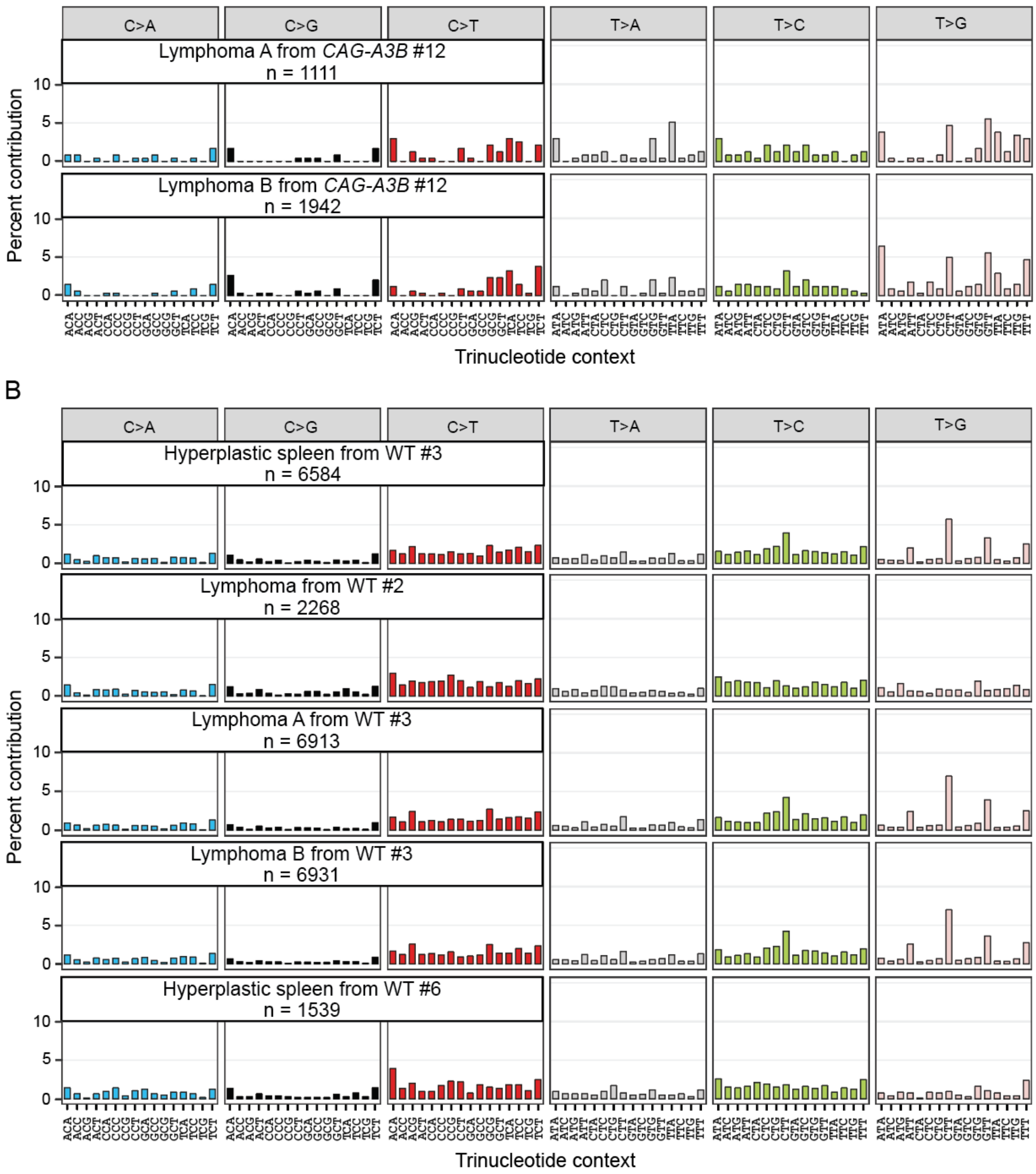


**Figure S9. Mutational spectra of blood tumors, related to Figure 5**

(**A-B**) Trinucleotide mutation profiles of all whole-genome sequenced blood tumors in *CAG-A3B* and WT mice not shown in Figure 5 (n-values = total SBS mutations in each tumor).

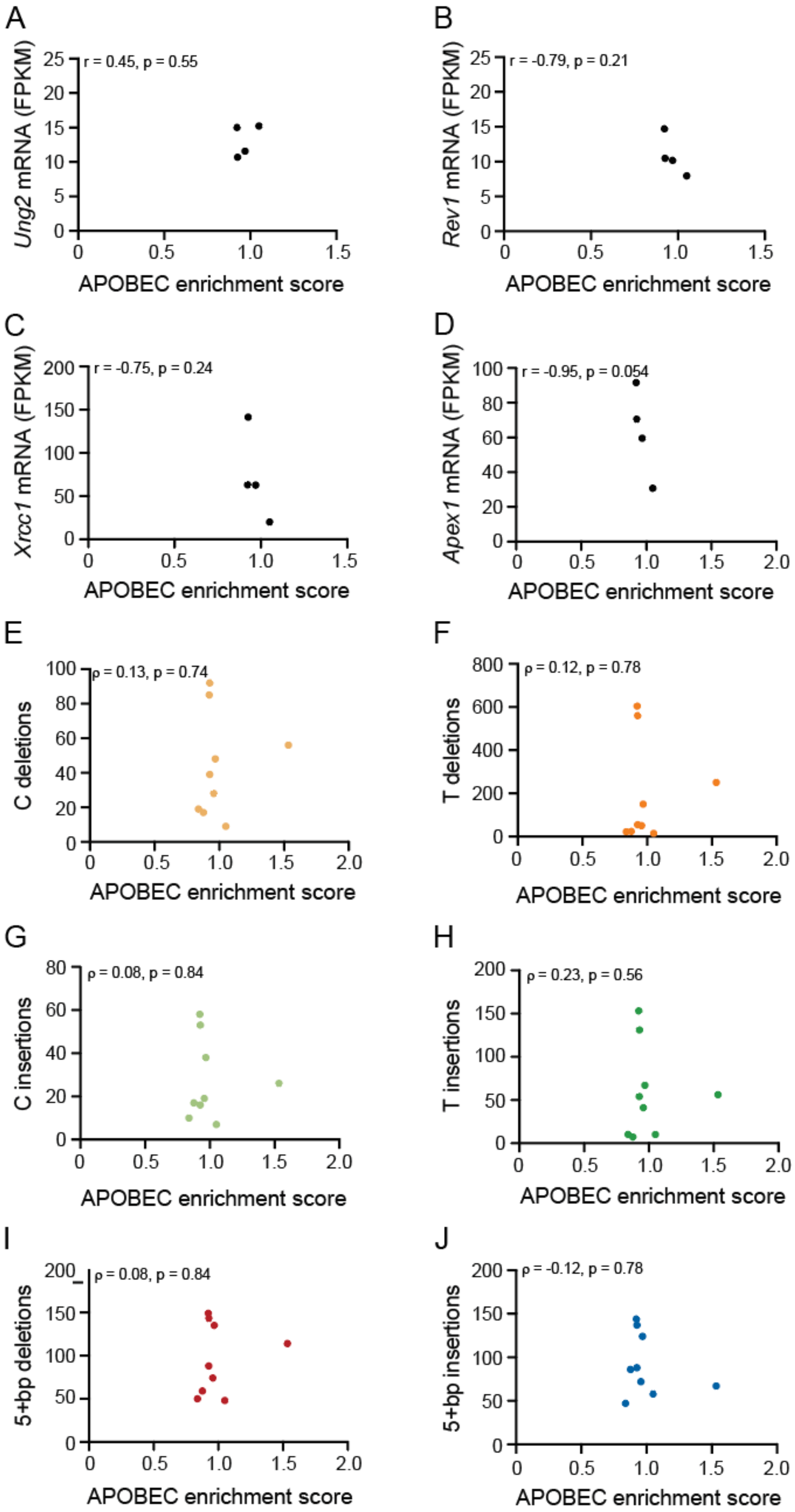
A
r = 0.45, p = 0.55
Ung2 mRNA (FPKM)
APOBEC enrichment score
B
r = -0.79, p = 0.21
Rev1 mRNA (FPKM)
APOBEC enrichment score
C
r = -0.75, p = 0.24
Xrcc1 mRNA (FPKM)
APOBEC enrichment score
D
r = -0.95, p = 0.054
Apex1 mRNA (FPKM)
APOBEC enrichment score
E
ρ = 0.13, p = 0.74
C deletions
APOBEC enrichment score
F
ρ = 0.12, p = 0.78
T deletions
APOBEC enrichment score
G
ρ = 0.08, p = 0.84
C insertions
APOBEC enrichment score
H
ρ = 0.23, p = 0.56
T insertions
APOBEC enrichment score
I
ρ = 0.08, p = 0.84
5+bp deletions
APOBEC enrichment score
J
ρ = -0.12, p = 0.78
5+bp insertions
APOBEC enrichment score

**Figure S10. Genomic correlations in WT mice, related to Figure 5**

(**A-D**) Scatterplots of APOBEC enrichment score from WT lymphomas (n=4) compared to the mRNA levels of *Ung2*, *Apex1*, *Xrcc1*, and *Rev1*, respectively, from the same tumors (Pearson correlation coefficients and corresponding p-values indicated).

(**E-J**) Scatterplots showing relationships between APOBEC enrichment scores from WT tumors and the indicated indel types (Spearman's rank correlation coefficients and corresponding p-values indicated).

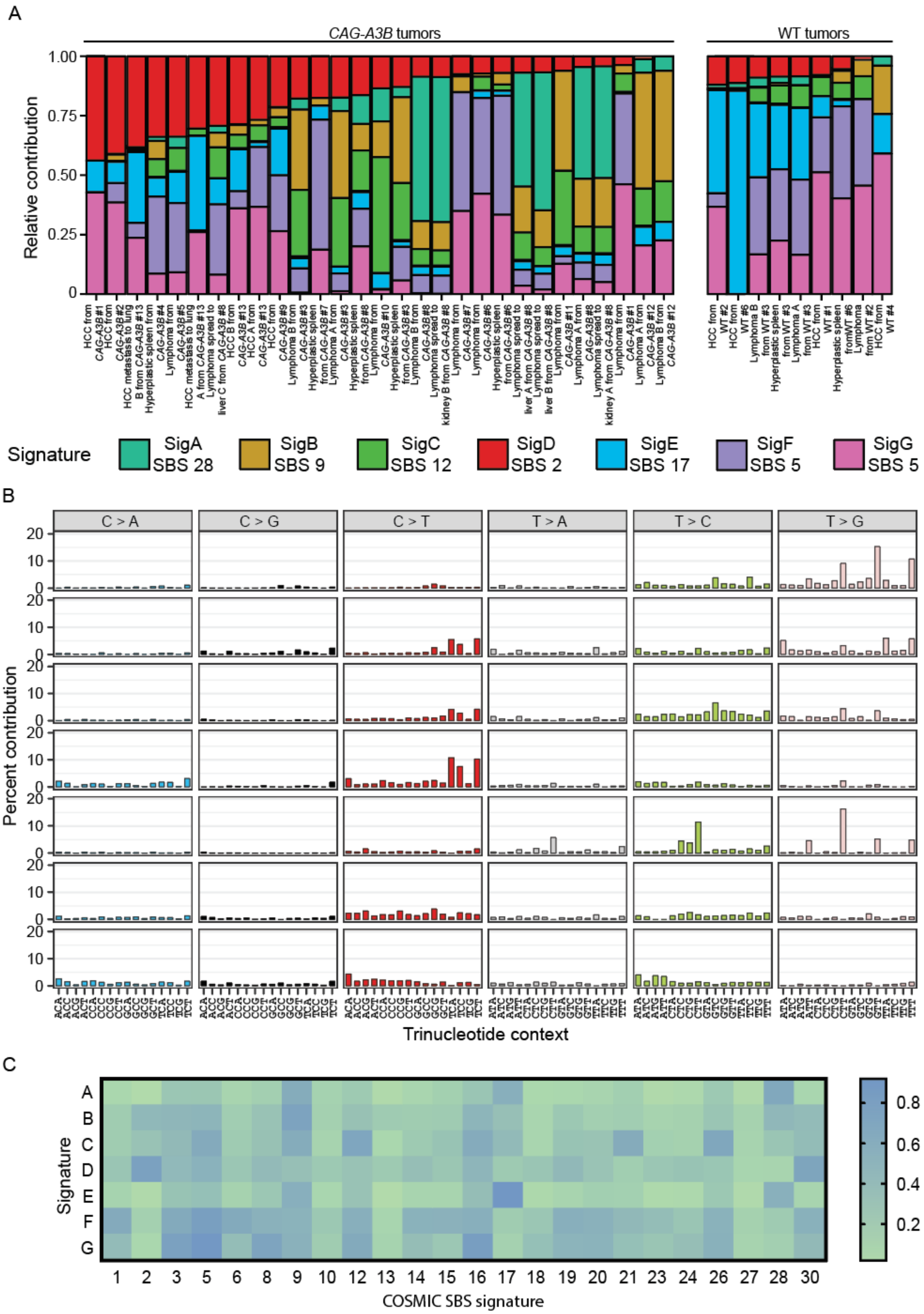
A
CAG-A3B tumors
WT tumors
Relative contribution
Signature
SigA SBS 28
SigB SBS 9
SigC SBS 12
SigD SBS 2
SigE SBS 17
SigF SBS 5
SigG SBS 5
B
C > A
C > G
C > T
T > A
T > C
T > G
Percent contribution
Trinucleotide context
C
Signature
COSMIC SBS signature

**Figure S11. Extraction of *de novo* signatures from all mouse tumors, related to Figure 5**

(**A**) Contribution of seven *de novo* signatures to the overall tumor mutation landscape based on non-negative matrix factorization (n=29 *CAG-A3B*; n=9 WT). Upon comparison to previously described COSMIC signatures, SigD is similar to SBS2 (cosine similarity, CS, = 0.82), SigA to SBS28 (CS = 0.72), SigB to SBS9 (CS = 0.78), SigC to SBS12 (CS =0.82), SigE to SBS17 (CS = 0.93), SigF to SBS5 (CS = 0.83), and SigG to SBS5 (CS = 0.92).

(**B**) Trinucleotide mutation profiles of *de novo* signatures SigA to SigG extracted from tumors using non-negative matrix factorization.

(**C**) Heat map showing cosine similarities of the 7 mutation signatures in the tumors described here in comparison to established COSMIC SBS mutation signatures. COSMIC SBS mutation signatures 4, 7, 11, 25, and 19 are excluded due to lack of relevance to mouse models.

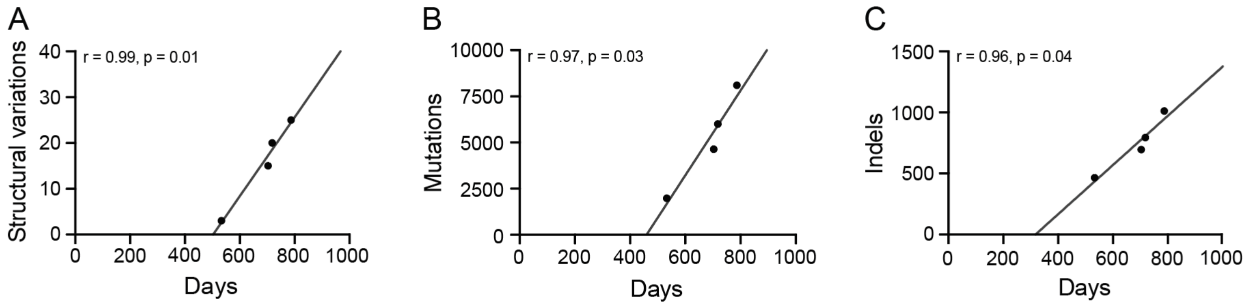


**Figure S12. Relationship between structural variations, SBS mutations, and indels and animal age, related to Figure 5 and 6**

(**A-C**) Scatterplot of the number of structural variations, SBS mutations, and indels in HCCs (n=4) from WT animals in comparison to age of the animal at the time of sacrifice (Pearson correlation coefficients and corresponding p-values shown). These positive associations indicate a linear relationship between mutation accumulation and age in tumors from WT animals.

**Table S1. Mice used in tumor-free survival and sequencing analyses**

Included as a separate file titled "Durfee-et-al-TableS1.xlsx".
